# Biophysical modeling reveals the transcriptional regulatory mechanism of Spo0A, the master regulator in starving *Bacillus subtilis*

**DOI:** 10.1101/2025.01.21.634141

**Authors:** Yujia Zhang, Cristina S.D. Palma, Zhuo Chen, Brenda Zarazúa-Osorio, Masaya Fujita, Oleg Igoshin

## Abstract

In starving *Bacillus subtilis* bacteria, the initiation of two survival programs – biofilm formation and sporulation – is both controlled by the same phosphorylated master regulator Spo0A∼P. Its gene, *spo0A*, is transcribed from two promoters, P*v* and P*s*, that are respectively regulated by RNAP holoenzymes bearing σ^A^ and σ^H^. Notably, transcription is directly autoregulated by Spo0A∼P binding sites known as 0A1, 0A2, and 0A3 box, located in between the two promoters. It remains unclear whether, at the onset of starvation, these boxes activate or repress *spo0A* expression, and whether the Spo0A∼P transcriptional feedback plays a role in the increase in *spo0A* expression. Based on the experimental data of the promoter activities under systematic perturbation of the promoter architecture, we developed a biophysical model of transcription regulation of *spo0A* by Spo0A∼P binding to each of the 0A boxes. The model predicts that Spo0A∼P binding to its boxes does not affect the RNA polymerase recruitment to the promoters but instead affects the transcriptional initiation rate. Moreover, the effects of Spo0A∼P binding to 0A boxes are mainly repressive and saturated early at the onset of starvation. Therefore, the increase in *spo0A* expression is mainly driven by the increase in RNAP holoenzyme levels. Additionally, we reveal that Spo0A∼P affinity to 0A boxes is strongest at 0A3 and weakest at 0A2 and that there are attractive forces between the occupied 0A boxes. Our findings, in addition to clarifying how the sporulation master regulator is controlled, offer a framework to predict regulatory outcomes of complex gene-regulatory mechanisms.

**Importance:** Cell differentiation is often critical for survival. In bacteria, differentiation decisions are controlled by transcriptional master regulators under transcriptional feedback control. Therefore, understanding how master regulators are transcriptionally regulated is required to understand differentiation. However, in many cases, underlying regulation is complex with multiple transcription factor binding sites and multiple promoters, making it challenging to dissect the exact mechanisms. Here, we address this problem for the *B. subtilis* master regulator Spo0A. Using a biophysical model, we quantitatively characterize the effect of individual transcription factor binding sites on each *spo0A* promoter. Further, the model allows to identify the specific transcription step that is affected by transcription factor binding. Such model is promising for the quantitative study of a wide range of master regulators involved in transcriptional feedback.

## 1. Introduction

To survive starvation conditions, *Bacillus subtilis* cells differentiate into two cell types. For mild starvation levels, cells can activate the production and secretion of an extracellular matrix, encasing a multicellular community called biofilm (1, 2). During prolonged starvation, cells can differentiate into metabolically inert spores that can survive long-term exposure to extreme environmental conditions (3–6).

The gene regulatory networks controlling biofilm and sporulation programs are triggered by the same master regulator Spo0A (0A) (4, 7–10), which is active in its phosphorylated form, 0A∼P (9, 11–13). Phosphorylation of 0A is controlled via a phosphotransfer cascade, termed phosphorelay (14, 7). The cascade efficiency regulates the post-translational activity of 0A (14, 7, 15, 10, 16, 17).

Meanwhile, 0A is also regulated at the transcriptional level. Specifically, expression of *spo0A* can initiate from two promoters, an upstream vegetative promoter (P*v*) and a downstream promoter (P*s*) (18, 19) (Figure 1, top panel). Each promoter is recognized by an RNA polymerase holoenzyme (RNAP) containing a specific sigma (σ) factors: housekeeping σ^*A*^ for P*v* and the alternative sigma factor σ^H^ for P*s* (19–22). In addition, transcription is directly autoregulated: four 0A∼P binding sites, known as 0A boxes, have been experimentally identified using DNase I footprinting assay (11, 23). Three of these are located downstream the P*v* promoter (0A1, 0A2, and 0A3). The fourth site, 0A4, overlaps with the -10 region of the P*s* promoter (23). These regulatory sites, together, form a transcriptional feedback loop and were shown to regulate *spo0A* expression (11, 23).

**Figure 1.**
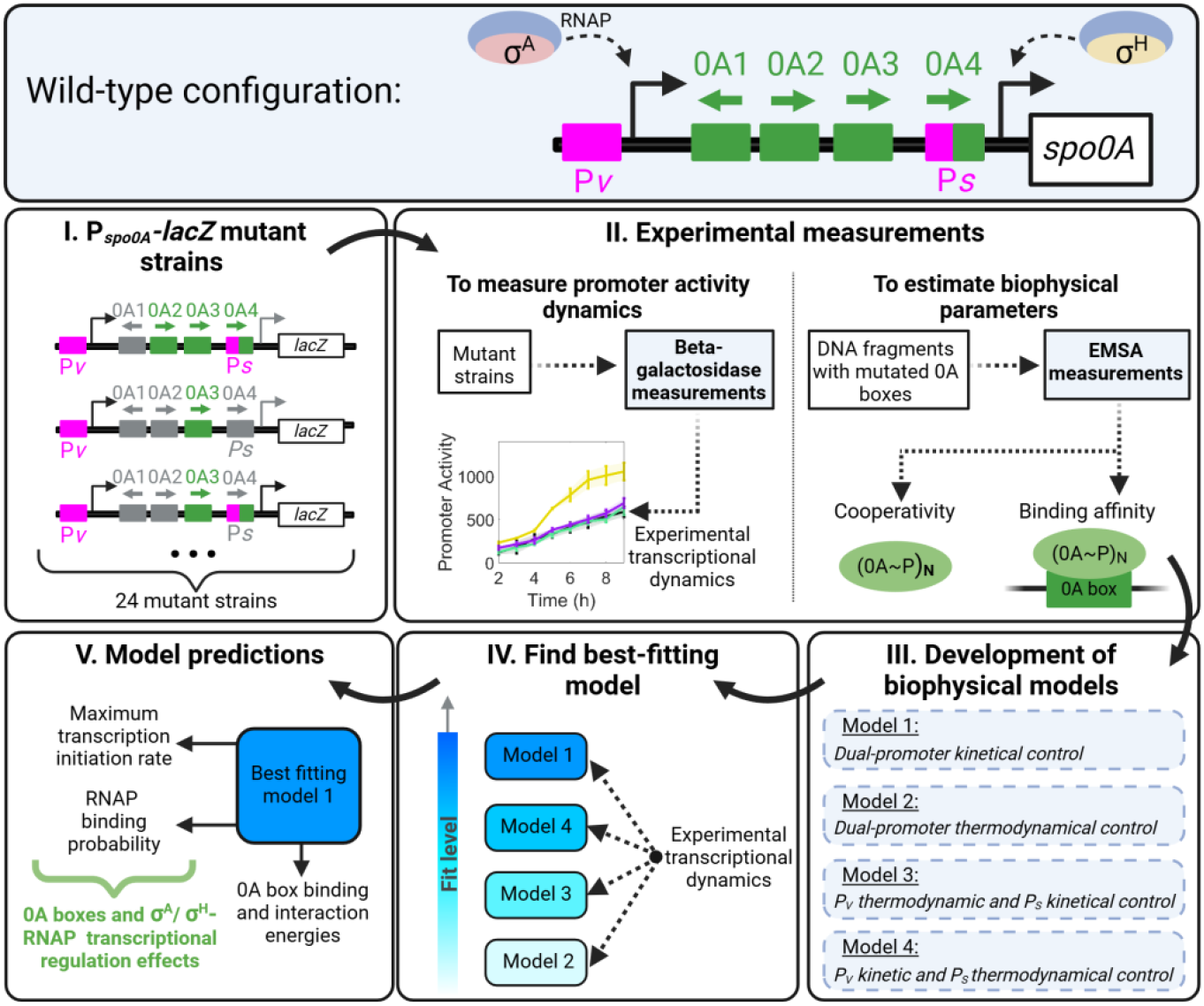
Study workflow. Top **panel:** Wild-type (WT) *spo0A* configuration. The gene *spo0A* is regulated by four transcription factor binding sites (0A boxes) and transcribed from two promoters (P*v* and P*s*). P*v* promoter is recognized by σ^A^-RNAP. P*s* promoter is recognized by σ^H^-RNAP. **Middle and lower panel: (I)** Types of P*spo0A-lacZ* mutant strains used in the study with all possible combinations of 0A boxes and promoter deletions. **(II)** Experimental β-galactosidase activity and electrophoresis mobility shift assay (EMSA) datasets published in (31). To estimate promoter activity, β-galactosidase activities were measured for each mutant strain. EMSA data was used to estimate 0A∼P binding affinity and cooperativity. **(III)** Development of all possible model combinations of promoter transcriptional control by considering four distinct scenarios. Model 1 assumes that the binding of 0A∼P has purely kinetic effects for P*v* and P*s* promoters. Model 2 assumes that the binding has purely thermodynamic effects for both promoters. Model 3 assumes binding has thermodynamic effects for P*v* and kinetic effects for P*s*. While in Model 4, the reverse occurs, with thermodynamic effects on P*s* and purely kinetic effects on P*v*. **(IV)** Using the EMSA estimated parameters, we find which model best fits the β-galactosidase activity data. **(V)** Study the model-predicted parameters to infer the regulatory role of RNAP and single or combinations of 0A boxes. Created with Biorender.com.

At the onset of starvation, *spo0A* expression increases (9, 11, 19, 20) concurrently with the increase in 0A∼P concentration and σ^H^ (10, 24). The increase in 0A∼P is a result of the increased cellular concentration of the upstream kinase activity due to the starvation-induced slowdown in cell growth rate (25, 6, 26, 27). The σ^H^ increases because 0A∼P represses the expression of *abrB*, encoding the transcriptional repressor of *sigH*, which encodes σ^H^. Thus, when 0A∼P level increases, the *sigH* repression is relieved, resulting in higher σ^H^ (therefore higher levels of σ^H^-RNAP) (28–30, 21, 24). As a result, *spo0A* is mostly transcribed from the P*v* promoter during vegetative growth, while P*s* promoter transcription is induced after the onset of starvation (18, 19). Notably, it remains unclear whether *spo0A* expression increases due to the transcriptional feedback. Moreover, it is yet to be determined whether the increase in one or both σ^A^- and σ^H^-RNAP levels is responsible for the increase in *spo0A* transcription.

To better understand *spo0A* autoregulation and pinpoint the exact role of 0A boxes upon 0A∼P binding, a recent study applied a systematic analysis of 0A box mutations and promoter elements, combined with biochemical assays for detecting interactions between Spo0A∼P and each 0A box (31). However, while the analysis of this data offered valuable insights, it did not yield quantitative conclusions. To address this, biophysical models, based on statistical mechanics, can integrate the accumulated datasets to predict the effect of transcription factor binding on the pattern of gene expression (32–37). Moreover, such models can independently examine the regulatory role of transcription factors and RNAP holoenzyme on gene expression.

Here, we develop a biophysical model for *spo0A* transcriptional regulation and use it to determine the regulatory roles of 0A boxes as well as σ^A^- and σ^H^-RNAP, at the onset of starvation. For this, we utilize a dataset (31) from strains with *lacZ* reporter expressed from genetically perturbed versions of *spo0A* promoter, including deleting one or multiple 0A boxes and/or deleting P*v* or P*s* promoter (Figure 1, step I). The dataset consists of *β*-galactosidase measurements of promoter activity for each strain. In addition, we also make use of electrophoretic mobility shift assay (EMSA) measurements (31) to estimate the 0A∼P cooperativity and binding affinity to each 0A box (Figure 1, step II). These data were then used in the biophysical models to study *spo0A* transcriptional regulation (Figure 1, step III). The models explored two different biophysical mechanisms (thermodynamic or kinetic) on how 0A boxes occupancy controls each promoter. We then compare the models in terms of their ability to best fit the data (Figure 1, step IV). In the end, we evaluate the model-predicted parameters to dissect the regulatory role of the 0A boxes and RNAP on the regulation of *spo0A* expression (Figure 1, step V). Overall, using mathematical modeling supported by experimental data, our study provides quantitative insights into the transcriptional autoregulation of the master regulator in the starving *B. subtilis*.

## 2. Result

### 2.1 *spo0A* promoter activity is regulated by 0A boxes

To understand the role of 0A boxes in regulating *spo0A* expression, we utilized promoter activity measurements of P*spo0A-lacZ* fusions to the wild type and 0A box genetically perturbed *spo0A* promoters (31), over an eight-hour period, beginning 2 hours after the start of culture in MSgg medium under biofilm and sporulation conditions (26, 27, 31). The perturbations included mutating (to eliminate 0A∼P binding; see Figure 1 of (31)) one or multiple 0A boxes and/or eliminating P*v* or P*s* promoter. Noteworthy, since the 0A4 box overlaps with the Ps promoter region, the elimination of the P*s* promoter removes the 0A4 box (23). For convenience, all *lacZ* fusion strains are hereafter categorized into three distinct promoter-specific groups, as shown in Figure 2A-C. The first group, referred to as ‘P*v*P*s* strains’ (Figure 2A), consists of strains that contain both promoters. The second group, termed ‘P*v* strains’, includes strains with only the P*v* promoter (i.e., *Δ*P*S*, Figure 2B). The third group corresponds to ‘P*s* strains*’* (i.e., *Δ*P*V*, Figure 2C). Each promoter-specific cohort is further composed of 8 strains, each of which carries single or multiple non-mutated 0A boxes (0A1, 0A2, and 0A3 and combinations thereof, abbreviated as ‘1’, ‘2’, ‘3’, …, and ‘123’). Each group also includes a strain with no 0A boxes (i.e., all 0A boxes were mutated), denoted as ‘none’. A complete description of the strains and their designations is provided in Supplementary Table S1 (31).

**Figure 2.**
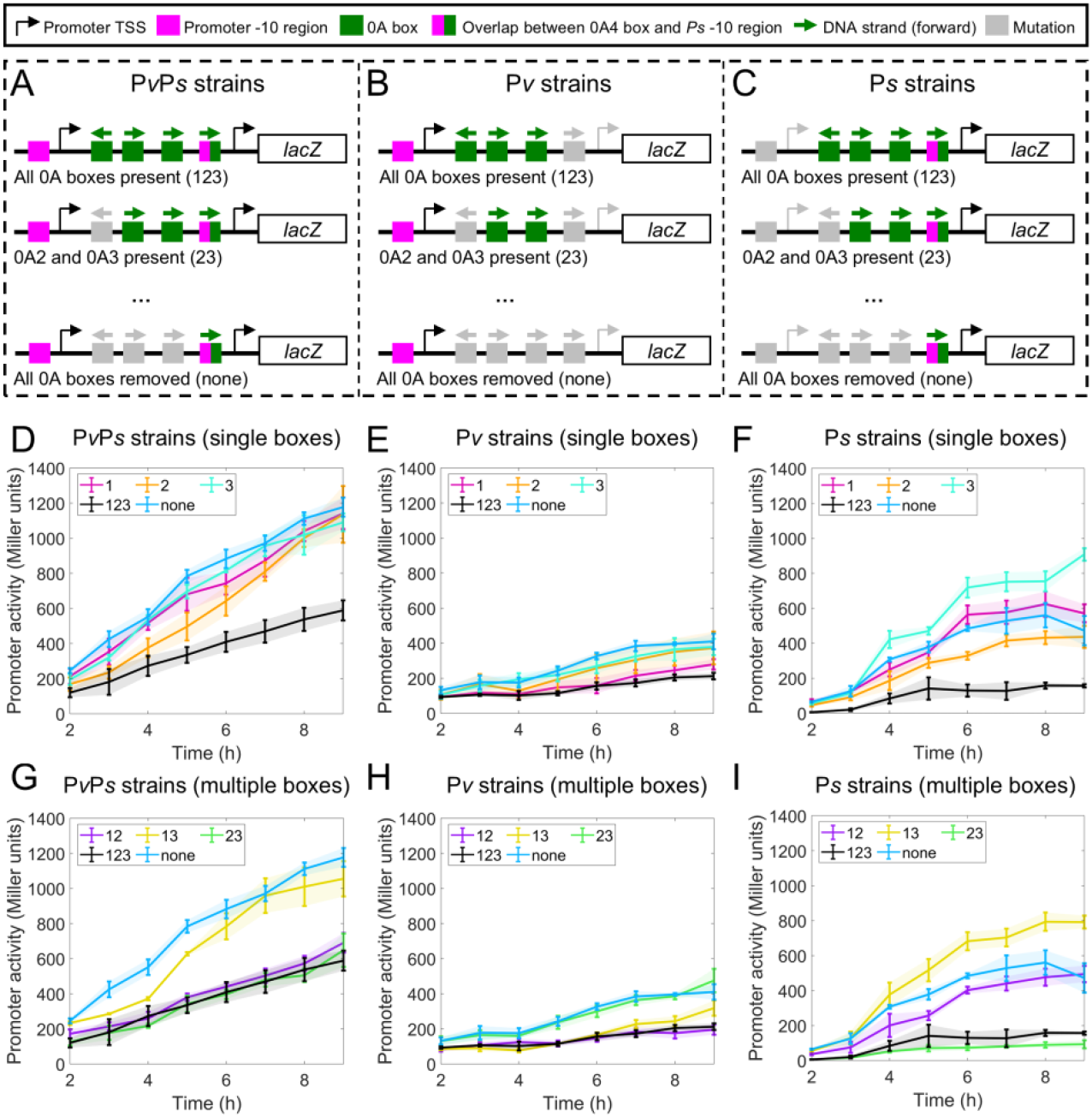
Experimental promoter activity measurements of *lacZ* fusion strains with different promoters and 0A boxes. **(A-C)** Illustration of the P*spo0A-lacZ* mutant strains constructed. Three cohorts of promoter specific constructs were built: **A)** no promoter deletion (‘P*v*P*s* strains’), **B)** P*s* promoter deletion (‘<u>P</u>*v* strains’), and **C)** P*v* promoter deletion (‘P*s* strains’). Note that the deletion of the P*s* promoter also removes 0A4 from ‘P*v* strains’. In addition, for the strains in each promoter-specific cohort, 0A1-3 boxes are not mutated at all, mutated one at a time, or mutated in combinations. The annotation of each strain is based on the presence of 0A1-3 boxes. For example, a strain with 0A1 (i.e., 0A2 and 0A3 mutated) is denoted as ‘1’ and a strain with all 0A1-3 boxes is denoted as ‘123’. For each promoter-specific cohort, a total of eight strains (all possible combinations of 0A box mutations) were constructed (not depicted in the figure). Complete strain details can be found in Supplementary Table S1 (31). **(D-F)** Experimentally measured promoter activity (β-galactosidase) over time for constructs with a single 0A box present (data from (31)). For comparison, in all plots, we also show the data for strains with all 0A boxes (‘123’, black lines) or no 0A boxes (‘none’, dark blue lines). **(G-I)** Experimentally measured promoter activity (β-galactosidase) over time for constructs with more than one 0A box present (data from (31)). For comparison, in all plots, we also show the data for strains with all 0A boxes (‘123’, black lines) or no 0A boxes (‘none’, dark blue lines). Vertical error bars are the standard deviation in biological replicates.

The promoter activity measurements demonstrate a complex pattern (Figure 2D-I). Namely, we observed that the effect of multiple 0A boxes cannot be explained by combining individual effects of single boxes. For instance, 0A∼P binding at 0A3 alone increases P*s* activity (Figure 2F, ‘3’ compared to ‘none’) and binding at 0A2 alone decreases P*s* activity (Figure 2F, ‘2’ compared to ‘none’). However, when 0A3 is simultaneously present with 0A2, P*s* activity is strongly repressed (Figure 2I, ‘23’ compared to ‘none’), with the repression being even more pronounced than that caused by 0A2 alone. Similar observations can be made for other strains. Overall, although the data is comprehensive, a simplistic qualitative analysis is insufficient to draw consistent conclusions about the regulatory consequences of 0A∼P binding across all genetic perturbations.

Despite the complexity, we found two ways to collapse the data. In the first approach, we normalize the data of each strain by its last timepoint measurement. Subsequently, each normalized strain measurement was further divided by the corresponding normalized measurement of the ‘none’ strain at each time point. The normalized data is shown in Figure 3A-F. The results show that, for most mutant strains, promoter activity data collapses to approximately one and remains relatively constant over time. While this data collapse shows a uniform trend, its interpretation remains unclear.

**Figure 3.**
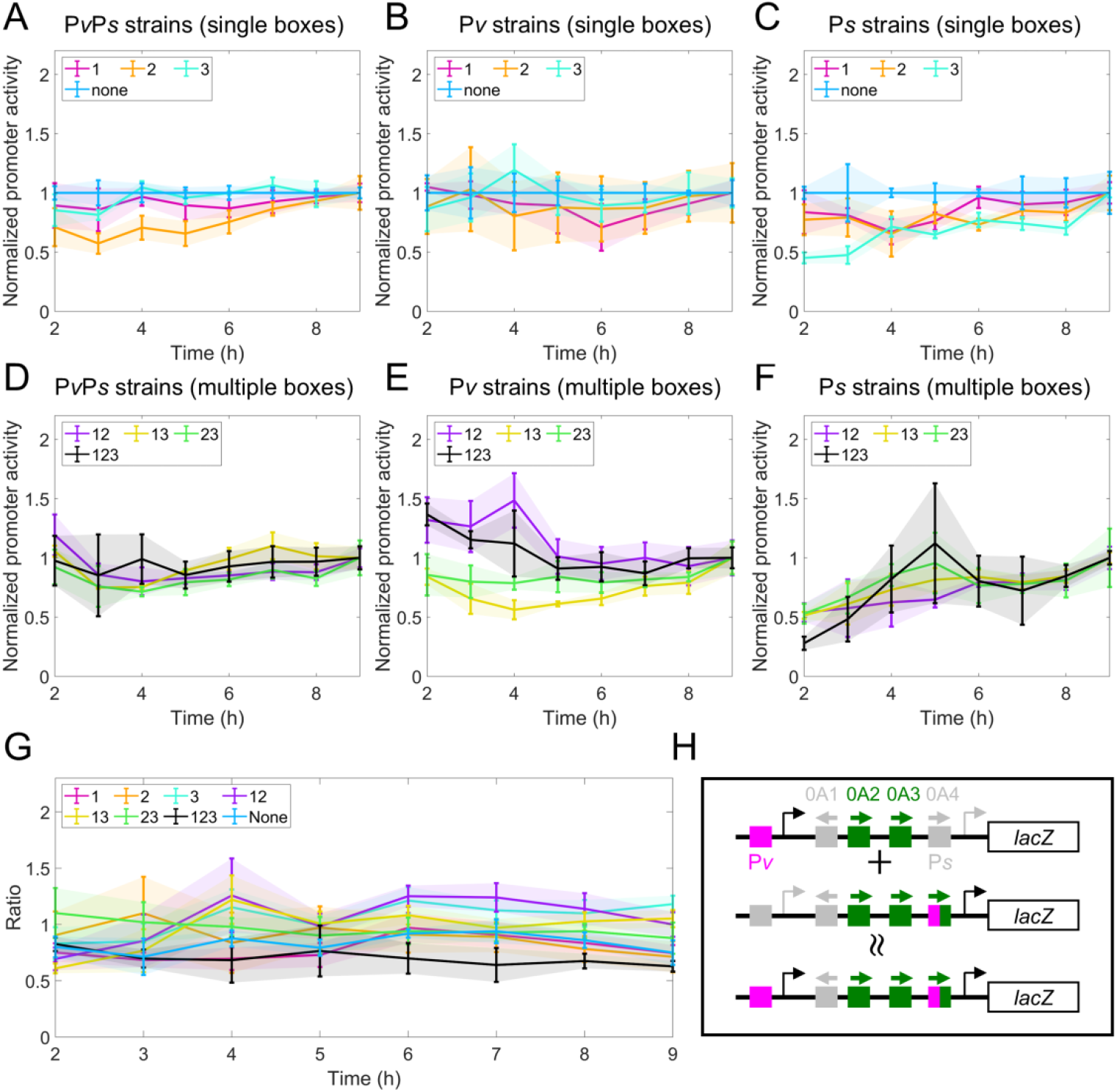
Data collapse of the *spo0A* promoter activity data. **(A-C)** Normalized promoter activity (β-galactosidase) over time for constructs with a single 0A box present, for each promoter-specific cohort. For comparison, in all plots, we also show the data for strains with no 0A boxes (‘none’, dark blue lines). **(D-F)** Normalized promoter activity (β-galactosidase) over time for constructs with more than one 0A box present, for each promoter-specific cohort. For comparison, in all plots, we also show the data for strains with all 0A boxes (‘123’, black lines). **(G)** Additivity level, i.e. the ratio computed by *Eqn. 1*, for all mutant strains over time. **(H)** Graphic illustration of what data collapse suggests about transcription of two promoters. Depicted is the sum of the promoter activity of ‘P*v* 23’ strain and ‘P*s* 23’ strain being approximately equal to the promoter activity of the ‘P*v*P*s* 23 strain’.

The second data collapse performed is to compute the ratio between promoter activity measurements of ‘P*v*<u>P</u>*s* strain’ and the sum of ‘<u>P</u>*v* strain’ and ‘P*s* strain’. For each mutant strain (*MS*) at each timepoint *t*, we compute the following ratio

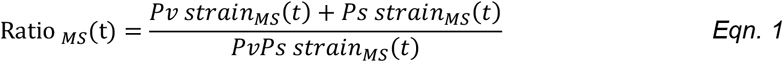

The results show that, at all times, for all mutant strains, the ratio is kept relatively constant and approximately equal to one (Figure 3G). In other words, the activity of the double promoters can be approximately estimated by adding together the single promoter activity, i.e., the single promoter expression is additive (see illustrative Figure 3H). Notably, this indicates that the transcription from each promoter happens independently of the other. Given the complexity in the experimental data and the data collapse of unclear origin, quantitative approaches are required to explain the transcriptional regulation of *spo0A*. Therefore, a mathematical model could provide valuable insights and a better understanding of the data.

### 2.2 0A boxes have different binding affinities to 0A∼P, and 0A∼P has tetrameric cooperativity

As a first step to constructing the mathematical model of gene regulation of the *spo0A* promoter, we need to determine how the occupancy of the different 0A boxes varies with increasing 0A∼P. To this end, we need to know the binding affinities of 0A∼P to the individual boxes. These data can be estimated from the measurements from gel electrophoresis mobility shift assay (EMSA) conducted for DNA fragments with systematic mutagenesis of 0A boxes, subject to various concentrations (0-2 *μ M*) of 0A∼P (31). For each 0A∼P concentration, the fraction of DNA fragments bound to 0A∼P was quantified. The experimental results for DNA fragments with a single 0A box present (Figure 4A, colored circles) show that, for increasing levels of 0A∼P, the fold increase in the fraction of bound DNA varies between strains. As such, these results suggest that individual 0A boxes have different binding affinities to 0A∼P.

**Figure 4.**
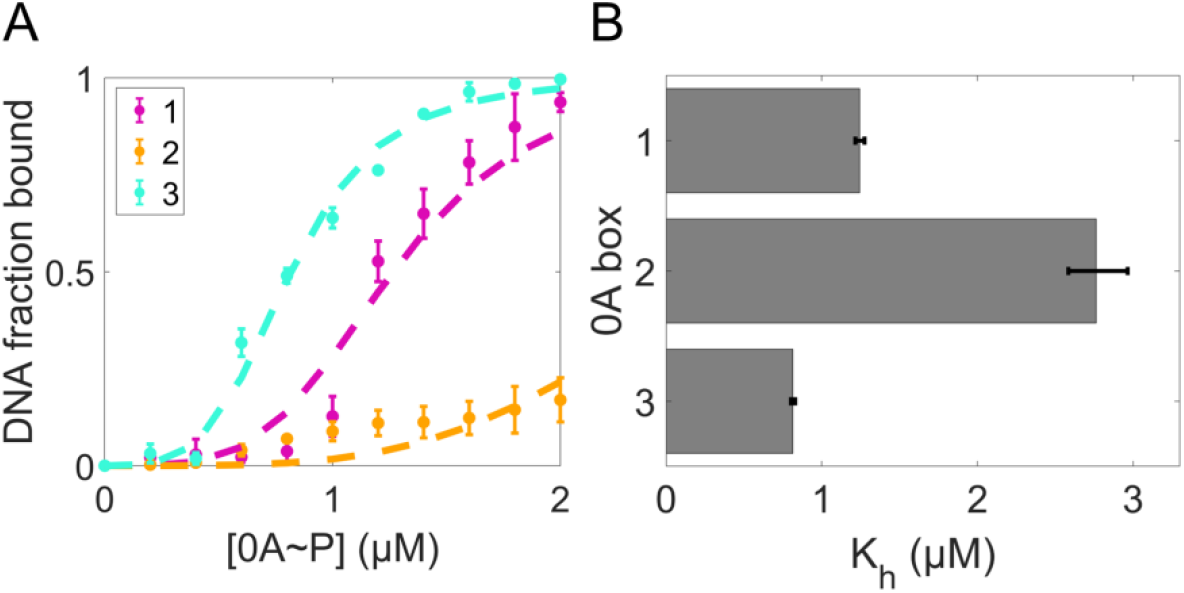
Estimate the number of 0A∼P bound to each 0A box and 0A box binding affinity *in vitro*. (**A)** The fraction of bound DNA under various 0A∼P concentrations, estimated from EMSA data (colored circles, data from (31)). The error bars represent standard deviation between biological duplicate experiments. The dashed lines are the best fitting results to *Eqn. 3*, assuming 0A∼P cooperativity (*N*) of 4. **(B)** Model-predicted concentration of 0A∼P at which the binding probability is half maximal (*K*_*h*_) for each 0A box. Error bars correspond to 95% confidence intervals estimated from 1000 fits to the augmented dataset.

To model the occupancy of each individual 0A boxes by 0A∼P and estimate binding affinities, we propose to use the following reaction scheme:

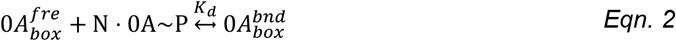

Here, *N* is the stoichiometric coefficient, i.e. cooperativity factor, that specifies the number of 0A∼P molecules that bind to a free 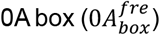 to form a bound 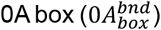, and *K*_*d*_ is the reaction dissociation rate constant. In accordance with the mass-action equilibrium, the fraction of 0A box bound to 0A∼P can be computed as a function of the number of 0A∼P molecules (*N*), 0A∼P concentration ([0*A*∼*P*]), and the concentration of 0A∼P at which the binding probability is half maximal (*K*_*h*_) (Supplementary Section I, subsection i.1):

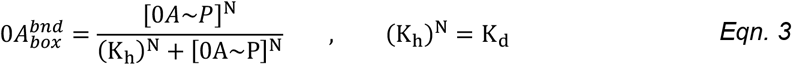

The magnitude of *K*_*h*_ is indicative of the 0A box binding affinity, i.e. it is the concentration of 0A∼P at which the box is occupied with 50% probability. Thus, a 0A box with lower *K*_*h*_ is more likely to be bound and has a high binding affinity.

To estimate *K*_*h*_ of individual 0A boxes, we fit *Eqn. 3* to the measured fraction of bound DNA for each DNA fragment with a single box, under known concentration of 0A∼P. To account for experimental uncertainties between replicates, an augmented data processing step (Supplementary section I, subsection i.2) was done prior to model fitting (Supplementary section I, subsection i.3). We started with assuming a dimer form of 0A∼P binding to the DNA (*N* = 2), as suggested by previous literature (38–40). However, this assumption failed to match the steepness of the curve in the experimental data as shown in Supplementary Figure S1. In contrast, *N* = 4 yields a good fit (Figure 4A, colored dashed lines), with an optimized error ∼3.7 fold lower than that when *N* = 2 (Supplementary Table S2).

Examination of the best-fit parameters allowed us to conclude that 0A3 box has the lowest *K*_*h*_, thus the highest binding affinity, followed by 0A1 and 0A2 (Figure 4B). In addition, based on *Eqn. 3*, we derived an equivalent statistical thermodynamical model to compute the fraction of bound DNA when more than a single 0A box is present (Supplementary Section I, subsection i.4). Using only the *K*_*h*_ parameters fitted from the single-box experimental measurements (Figure 4B), the model produced predictions that explain well the EMSA measurements with multiple 0A boxes (Supplementary Figure S2). Such minimal model, without additional parameters, suggest the intermolecular interactions between 0A∼P molecules are not significant under *in vitro* experimental conditions.

### 2.3 The purely kinetic control model best fits *spo0A* promoter activity, is consistent with experimental data in terms of additivity, and predicts 0A∼P interaction energies

To determine the mechanisms of transcriptional regulation of 0A boxes on *spo0A* promoters, we developed a biophysical model of gene regulation to predict the transcriptional rate (Figure 2D-I). In accordance with previous literature (32–34, 36, 37), we postulate that the effective transcription rate (*v*^*eff*^) is computed as the weighted sum of the probability of RNAP being bound to the promoter (*P*) and the maximum transcription initiation rate (*v*_*max*_), over all promoter bound configurations, *α* ∈ *α*_*T*_ :

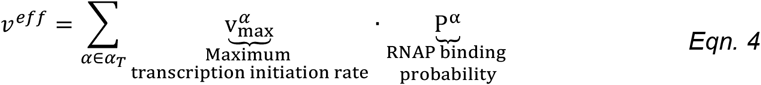

Detailed information on how to compute the probabilities and the maximum transcription rate as a function of energy parameters is described in Materials and Methods, Section 4.4 and Supplementary Sections II and III. According to the transcriptional function (*Eqn. 4*), gene expression can be regulated by affecting the RNAP binding probability (thermodynamic control) and by affecting the maximum transcription initiation rate (kinetic control). For purely thermodynamic control, it is usually assumed that the transcription initiation rates are the same and the RNAP binding probability is affected by attractive or repulsive interaction energies between bound 0A∼P to each 0A box and RNAP (Figure 5A, RNAP binding reaction affected). For purely kinetic control, it is assumed interaction energies do not exist, but transcription initiation rates differ (Figure 5A, transcription initiation reaction affected) depending on the binding configuration *α* (binding configurations considered are in Supplementary Table S3).

**Figure 5:**
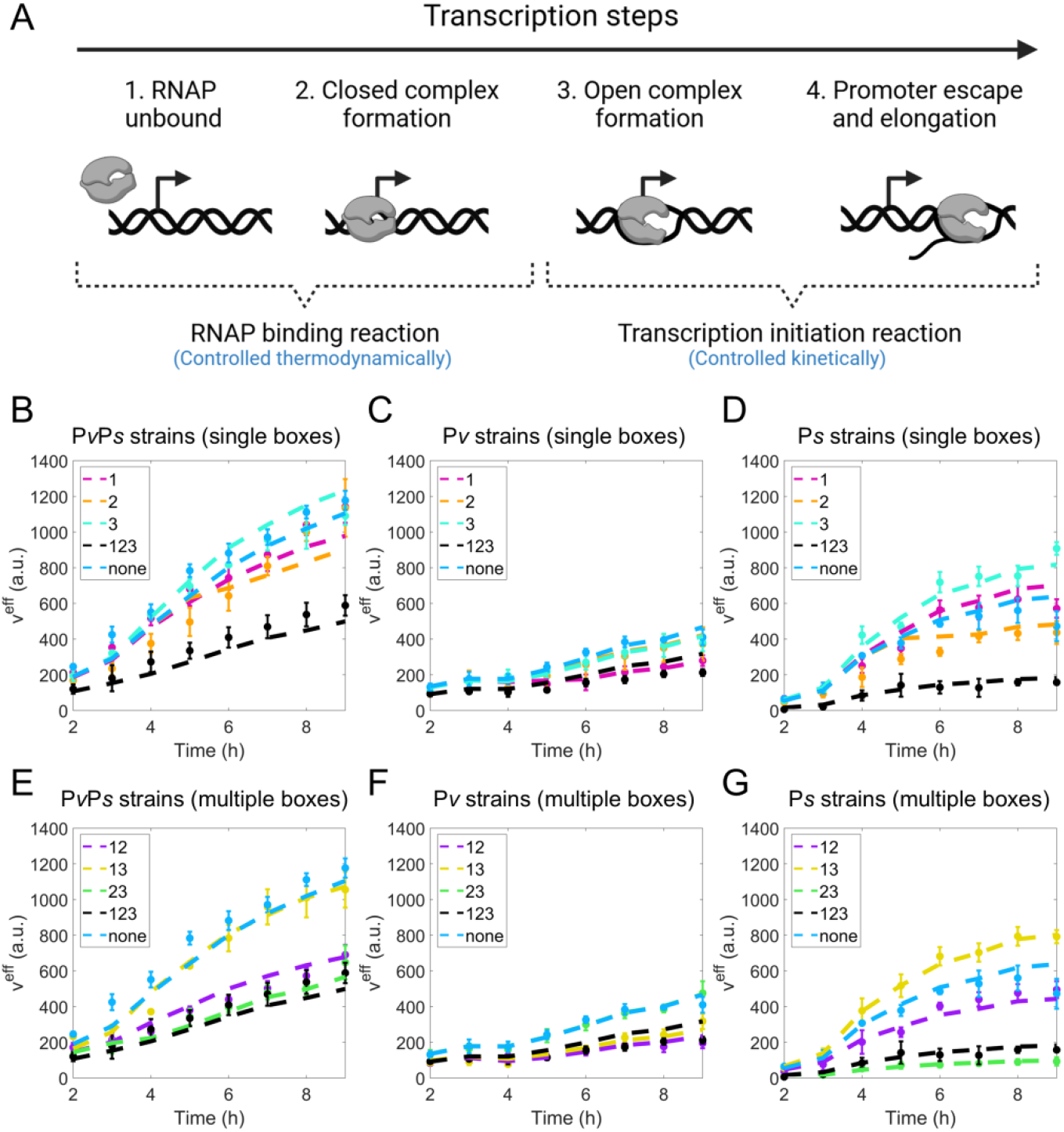
A purely kinetic control model best explains the experimental measurement of *spo0A* promoter activity. **(A)** Schematic illustration of transcription steps. Shown from left to right are the postulated transcription steps: First, RNAP recognizes and binds to the promoter forming the closed complex. This reaction rate is controlled thermodynamically and governed by the RNAP binding probability (*P*). Next, transcription initiation reaction begins with the closed complex isomerizing into an open complex (41), followed by RNAP promoter escape and transcription elongation (42, 43). The transcription initiation reaction is governed by the maximum transcription initiation rate (*v*_*max*_) and controlled kinetically. The model regulates the effective transcription rate (*v*^*eff*^) by affecting the RNAP binding probability (*P*) and by affecting the transcription initiation rate (*v*_*max*_). **(B-D)** Shown are the purely kinetic model fitting results (*v*^*eff*^) for strains with the presence of a single box in ‘P*v*P*s* strains’, ‘P*v* strains’, and ‘P*s* strains’, respectively. **(E-G)** Shown are the purely kinetic model fitting results (*v*^*eff*^) for strains with the presence of multiple 0A boxes for the ‘P*v*P*s* strains’, ‘P*v* strains’, and ‘P*s* strains’, respectively. For all plots, strains with no 0A boxes (‘none’, dark blue) and strains with all 0A boxes (‘123’, black) are also shown for comparison. For comparison, the experimental time-dependent promoter activity data for each mutant strain is also shown (data points with error bars). Error bars are the standard deviation of more than two biological replicates.

Here, we consider four model scenarios since there are two promoters regulating *spo0A* and each promoter can follow one of the two biophysical mechanisms. In the first scenario, the binding of 0A∼P has purely thermodynamic effects for both P*v* and P*s* promoters (Supplementary Section IV, subsection iv.1). In the second, the binding has purely kinetic effects for both P*v* and P*s* promoters (Supplementary Section IV, subsection iv.2). The third scenario assumes purely thermodynamic effects for P*v* and purely kinetic for P*v*. While in the fourth scenario, the reverse occurs, with thermodynamic effects on P*s* and purely kinetic effects on P*v* (Supplementary Sections IV, subsection iv.3).

Noteworthy, the model focuses on the regulatory roles of 0A1–0A3 boxes, excluding 0A4, based on the observations in the experimental promoter dynamics. First, the additive property observed in promoter activity measurements (Results section 2.1) suggests that the 0A4 box has a negligible regulatory impact on the P*v* promoter. Specifically, the combined activity of P*v* strains (lacking the 0A4 box) and P*s* strains (which include the 0A4 box) matches the activity of P*s*P*v* strains (which include the 0A4 box). Second, the absence of repression in P*s* strains, even at later time points (Figure 2F and 2I), implies that 0A∼P binding to the 0A4 box is unlikely to prevent σ^H^-RNAP binding, despite the overlap between the 0A4 box and the P*s* promoter region (11, 23). Overall, we find 0A4 to have minimal effects in P*s* and P*v* promoters.

To predict the time-dependent activation of gene expression as shown in experimental data (Figure 2D-I), we need time-dependent concentrations of the transcription factor (0A∼P) and RNAP holoenzyme. We assumed the timely increase in 0A∼P concentration to follow the previously predicted dynamics (26), also shown in Supplementary Figure S3. The σ^H^-RNAP and σ^A^-RNAP levels were kept as fitting parameters. The assumption of dynamic σ^A^-RNAP levels is supported by the experimental data showing that the activity of the ‘P*v* strain, none’ increases as a function of time (Figure 2E). In addition, increasing levels of σ^H^-RNAP is consistent with the existing literature (28, 24, 10).

With the above assumptions, we then fit the four models to the time-dependent P*spo0A-lacZ* measurements, respectively (Materials and Methods, Section 4.5), constrained by the half saturation concentration of 0A boxes (Figure 4B) and 0A∼P cooperativity estimated from EMSA measurements (Results section 2.2) (see Supplementary Section V for implementation of constraints). The model with purely kinetic control for both promoters was found to have the lowest optimization error (Supplementary Figure S4), hence it best fits the experimental data. Therefore, we conclude that 0A∼P binding to 0A boxes, in both P*v* and P*s* promoter, does not affect the RNAP binding (recruitment) to the promoter. Instead, the presence of 0A∼P at the promoter modulates the maximum transcription initiation rate (Figure 5A). The model results for each mutant strain and the experimental data used for fitting are shown in Figure 5B-G. All model-predicted parameters are shown in Supplementary Table S4.

Notably, our prediction of the purely kinetic control for both promoters can explain the additivity data collapse (Figure 3G, Results section 2.1). A model with purely kinetic control always leads to an additive transcription rate between two promoters; this can be demonstrated analytically (Supplementary Section IV, subsection iv.2.b). On the contrary, purely thermodynamically controlled promoter expression is generally not additive (Supplementary Figure S5). This can be intuitively understood by considering a DNA fragment with one 0A box, P*v*, and P*s* promoter. Under thermodynamic control, the bound 0A∼P interacts with both the bound RNAP at P*v* and at P*s*. As such, the activity from the two promoters is not independent, causing individual promoter activity to not be additive.

### 2.4 0A box effects are mainly repressive and saturated by the onset of starvation

The best-fit kinetic model allows to examine the mechanism of *spo0A* autoregulation. To this end, we first analyzed how occupancy of the different 0A boxes varies with increasing 0A∼P level. (Supplementary Figure S6 and Supplementary Section VI). We found that, for ‘123’ strains (i.e., all boxes present), all 0A boxes are already saturated by the onset of starvation (i.e. t = 2 h). All other strains reach 0A box saturation at varying timings, but all being saturated by t = 9 h. The early saturation of 0A∼P binding for the ‘123’ strain, even at low levels of 0A∼P, is due to the attractive interactions between bound 0A boxes. Specifically, the best-fit model predicts that the interaction energy between 0A2 and 0A3 boxes is the most attractive, while the interaction energy between 0A1 and 0A3 is comparable to that between 0A1 and 0A2 (Supplementary Table S4). To confirm that the predicted attractions between bound 0A boxes are necessary for the regulation of *spo0A* expression, we examined whether the model could fit well in the absence of one or more types of 0A box interaction, i.e., when interaction energy fitting parameters are constrained to be zero (Supplementary Table S5). Our results show that the model can only provide a good fit with the existence of all secondary interaction energies between 0A boxes. We further confirmed that the addition of such interaction energy parameters did not lead to model overfit by comparing Bayesian Information Criterion (41) (BIC) of the differentially constrained kinetic control models (Supplementary Section VII).

We then examined the effects of 0A∼P binding at individual 0A boxes. To this end, we started by analyzing the change in the model-predicted maximum transcription initiation rate for each 0A box-bound configuration (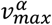in *Eqn. 4*). If a 0A box has a repressive regulatory effect, a bound 0A box will lead to a lower *v*_*max*_. The *v*_*max*_ of each possible 0A∼P bound configuration relative to the unbound 0A box configuration (‘000’), for each strain, is shown in Figure 6A-C. Overall, *v*_*max*_ < 1 for ‘P*v* strains’ and ‘P*v*P*s* strain’. As such, for these strains, the binding of 0A∼P mostly reduces *v*_*max*_, whether individual or combinations of 0A boxes are bound. Moreover, 0A∼P binding is also mostly repressive for P*s* as evidenced by *v*_*max*_ < 1 in ‘P*s* strains’, even though some 0A boxes such as 0A1, 0A3, and the combination thereof, have activating effects.

**Figure 6.**
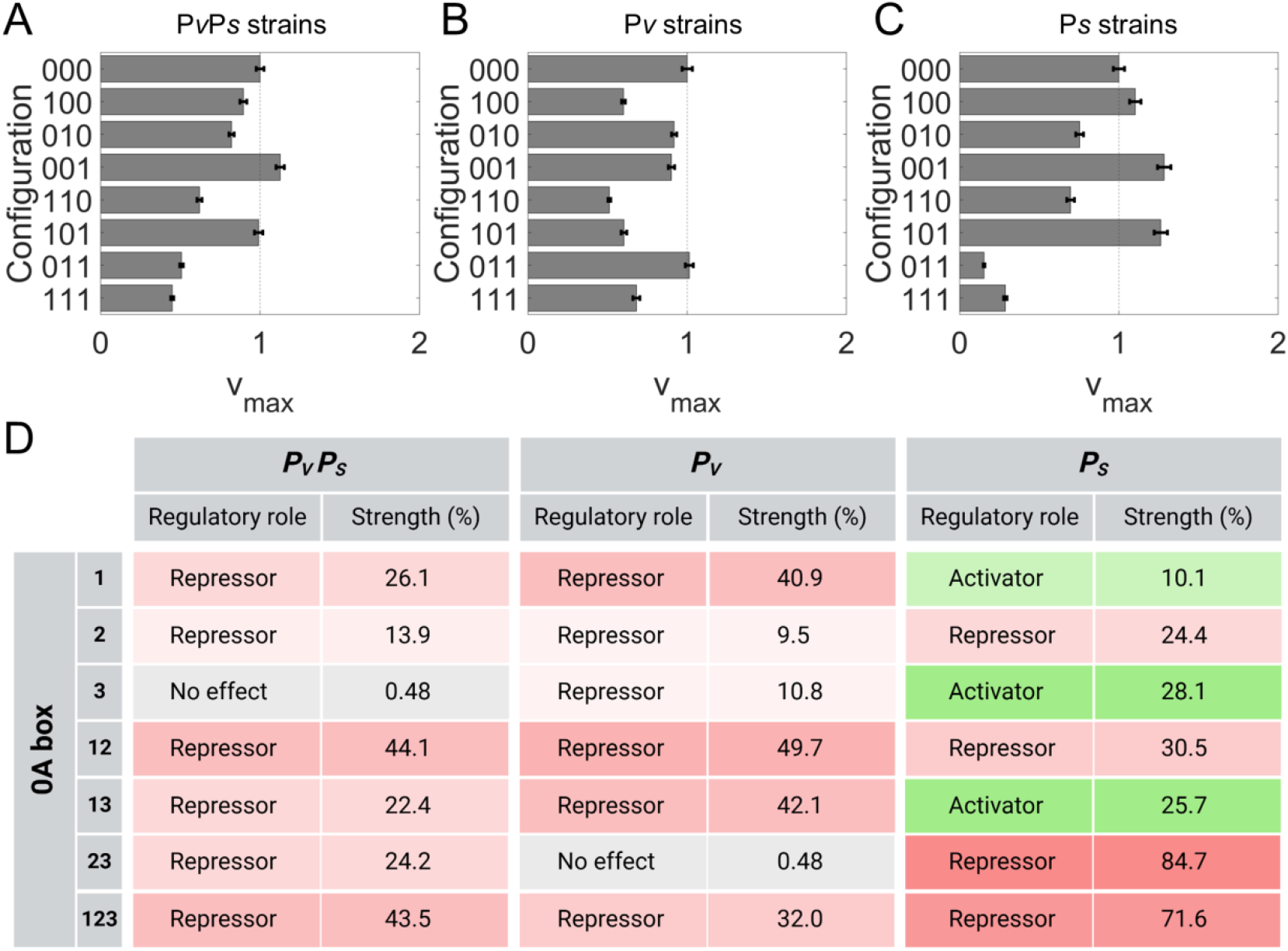
Role of 0A∼P binding in regulation of *spo0A* expression. **(A-C)** All model-predicted *v*_*max*_ parameter for each configuration relative to the configuration with no 0A box bound to 0A∼P ‘000’. Each plot corresponds to a promoter-specific construct, ‘P*v*P*s* strains*’*, ‘P*v* strains’, and ‘P*s* strains’, respectively. The strain configuration is represented by a binary sequence of three digits. Each digit, from left to right, corresponds to 0A1, 0A2, and 0A3 box, respectively. A value of “1” denotes a bound 0A box while “0” denotes unbound. **(D)** Effect of the presence of single and combinations of 0A boxes on the dual promoter activity (P*v*P*s*), in P*v* promoter, and in P*s* promoter. The repressive or activation strength is the percent increase or decrease of the expression of a mutant strain with respect to the expression of the ‘none’ strain (no 0A boxes at all), at t = 9h, assuming only 0A∼P concentrations are time-dependent (Supplementary Figure S7). Gradient colors represent the strength intensity, red for repression and green for activation.

In addition to examining *v*_*max*_, we also analyzed the effect of 0A∼P binding on the effective transcription rate (*v*^*eff*^). For this, we computed *v*^*eff*^ with 0A∼P concentration being time-dependent but RNAP concentration being constant over time (equal to t = 2 h for all times). This ensures that variations in expression levels over time result solely from 0A box control. We then measured the percentage increase or decrease in the model-predicted expression of each strain relative to the expression of the ‘none’ strain (strain with all 0A boxes mutated) at t = 9 h (Supplementary Figure S7). The results are shown in Figure 6D. Specifically, 0A1 represses P*v* more than 0A2 and 0A3. Meanwhile, for P*s*, both 0A1 and 0A3 are activators, whereas 0A2 is a repressor. For the dual-promoter strains, 0A3 shows negligible effects, and 0A2 is slightly less repressive than 0A1. The presence of multiple boxes holds different effects. For example, for P*s*, the simultaneous presence of 0A2 and 0A3 show repressive effects, overriding the single 0A3 box activation effects. Noteworthy, since 0A∼P binding is saturated by t = 9 h for all mutant strains, for ‘P*v’* and ‘P*s’* strains, the relative change in *v*_*max*_ (Figure 6A-C) and *v*^*eff*^ (Figure 6D) is the same. For example, a P*s* strain has a 71.5 % lower *v*_*max*_ when 0A1-3 is bound (Figure 6C, ‘111’ compared to ‘000’). Consistently, *v*^*eff*^ of ‘P*s* strain, 123’ is also 71.5 % lower (Figure 6D) than ‘P*s* strain, none’. On the other hand, for ‘P*v*P*s*’ strains, this trend does not exist. Instead, the change in *v*^*eff*^ is expected to be a weighted sum of the change in *v*_*max*_ of ‘P*v’* and ‘P*s’* strains (Supplementary Section VIII), i.e., at 0A∼P binding saturation, the regulatory effects of 0A∼P binding on P*v*P*s* is somewhere in between the regulatory effects on P*v* and P*s* alone. Overall, given that the WT strain (i.e. ‘P*v*P*s* 123 strain’) shows a 43.5 % repression strength and 0A box saturation by t = 2 h, we conclude that the effect of 0A boxes on *spo0A* transcription is mostly repressive and saturated by the onset of starvation. As such, a separate activating mechanism, other than 0A∼P binding, is responsible for the increase in *spo0A* expression after the onset of starvation (Figure 2A-F).

### 2.5 Increases in σ^A^- and σ^H^-RNAP levels regulate the increase in *spo0A* expression after the onset of starvation

The best-fitting model predicts that the effective transcription rate of strains with no 0A boxes (‘none’ in Figure 7A-C) increases over time, for ‘P*v*P*s*’, ‘P*v*’ and ‘P*s*’ strains. Since no 0A∼P binding occurs in these strains, such increase can only be explained through the time-dependent changes in RNAP holoenzymes. Therefore, our model predicts that RNAP holoenzyme dynamics (i.e. σ^A^- and σ^H^-RNAP) upregulate the *spo0A* expression over time, after the onset of starvation.

**Figure 7.**
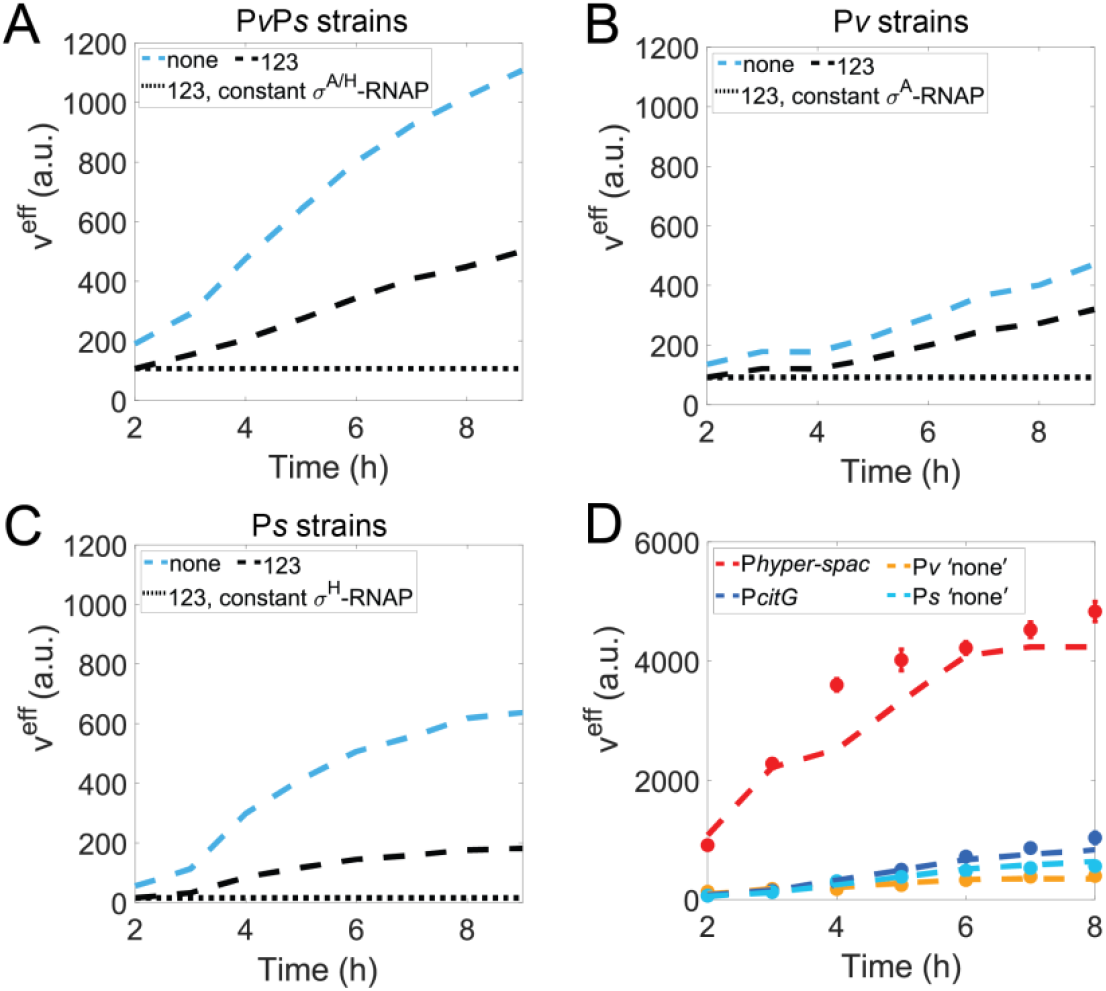
Role σ^A^- and σ^H^-RNAP in the regulation of *spo0A* expression. Model predicted effective transcription rate (*v*^*eff*^) assuming: **(A)** constant *σ*^*A*^- and constant *σ*^*H*^-RNAP levels for ‘P*v*P*s* strains, 123’; **(B)** constant *σ*^*A*^- RNAP levels for ‘P*v* strains, 123’; **(C)** constant *σ*^*H*^- RNAP levels for ‘P*s* strains, 123*’*. In panels **(A-C)**, also shown are the model-predicted dynamics for the two reference strains: the ‘none’ (i.e. strain with no 0A boxes) and the ‘123’ strain (i.e. strain with all WT 0A boxes), both of which are computed with dynamical RNAP levels. **(D)** Promoter activity measurements of P*hyper-spac*, P*citG*, P*v ‘*none’ (i.e. no 0A boxes present) and P*s ‘*none’ (shown in filled circles). Vertical error bars are standard deviation of at least two biological replicates. Also shown are the models predicted dynamics (dashed lines) assuming P*hyper-spac* and P*v*, P*citG* and P*s* have different RNAP binding affinities, but same σ^A^- and σ^H^-RNAP dynamics, respectively.

To illustrate this further, we used our best-fit model to compute the *spo0A* effective transcription rate (*v*^*eff*^) for the strains with all 0A boxes, using either a constant level (black dotted lines in Figures 7A-C,) or the model-predicted σ^A^- and σ^H^-RNAP dynamics (black dashed lines in Figure 7A-C). The constant level was set to equal to the model-predicted dynamics at t = 2h for all times (see Supplementary Figure S8 for model-predicted σ^A^- and σ^H^-RNAP dynamics). We found that, for the single promoter constructs, assuming constant levels of the respective RNAP (σ^A^ for ‘P*v* strain’ and σ^H^ for *‘*P*s* strain’) leads to an almost negligible increase in *spo0A* expression (black dotted lines in Figures 7B and 7C). In addition, for the dual promoter construct, assuming constant levels of both σ^A^- and σ^H^-RNAP predicts almost constant expression for *spo0A* (black dotted lines in Figure 7A), which is expected given the additivity property of the results. Altogether, these results suggest that σ^A^- and σ^H^-RNAP increase is the predominant regulatory mechanism responsible for *spo0A* activation at the onset of starvation.

Notably, this finding can explain the first data collapse results shown in (Result Section 2.1, Figure 3A-F). During data processing, the normalization of measurements by the ‘none’ strain measurements effectively removes the effect of RNAP binding. Consequently, due to a lack of activating regulatory mechanism, promoter activity remains relatively constant over time, evidenced by the collapse centered around 1 upon further normalization by the final time point. When we perform the same data collapse on the model-predicted effective transcription rate (Figure 5B-G), the results demonstrate a similar pattern for all mutant strains (Supplementary Figure S9). As such, our model predicts that, in all mutant strains, the RNAP dynamics is responsible for activating *spo0A* expression after the onset of starvation. In Supplementary Table S6, we dissect the contribution of RNAP holoenzyme dynamics from the effects of 0A∼P binding on *spo0A* expression, for each strain, after the onset of starvation.

We note that our prediction of increased σ^A^ and σ^H^ activities should not be limited to spo0A promoter. Therefore, to experimentally validate the model prediction that σ^A^- and σ^H^-RNAP is time-dependent, we measured the promoter activity of other σ^A^ and σ^H^ dependent promoters (strain description in Supplementary Table S1). Specifically, constitutive promoter activity was measured for *hyper-spac* promoter (P*hyper-spac*, σ^A^ specific) (9, 42) and *citG* promoter (P*citG*, σ^H^ specific) (43). The results (Figure 7D, colored data points for ‘P*hyper-spac’ and ‘*P*citG’*) show that the activity of both promoters increases over time in the absence of additional regulatory mechanisms such as transcription factor binding, further supporting the crucial role of both σ^A^- and σ^H^-RNAP activity in upregulating gene expression, since the increase in activity is not limited to the transcription of *spo0A*.

Furthermore, we hypothesized that the observed increase in promoter activity of P*hyper-spac* and P*v*, as well as P*citG* and P*s*, can be explained by the same levels of the σ^A^-and σ^H^-RNAP, respectively. To test for this, we examined if the biophysical model that assumes the same σ^A^-RNAP dynamics but different RNAP binding affinity (to account for different promoter binding affinities, as evidenced in Supplementary Figure S10) could successfully fit the P*hyper-spac* and *‘*P*v* strain none’ experimental measurements simultaneously. We find the model to fit both well (Supplementary Section IX) (dashed navy and light blue lines in Figure 7D). The same is found to be true for P*citG and* ‘P*s* strain none’, when assuming the same σ^H^-RNAP dynamics (dashed red and orange lines in Figure 7D). As such, we conclude that the biophysical model correctly predicted σ^A^- and σ^H^-RNAP increasing dynamics and that these are the predominant regulatory mechanisms responsible for the upregulation of *spo0A* over time, after the onset of starvation.

## 3. Discussion

The expression of *spo0A* is regulated by two promoters (P*v* and P*s*) with different RNAP holoenzyme preferences (bearing σ^A^ and σ^H^, respectively) (22) and by three regulatory regions located in between the promoters, designated as 0A1, 0A2, and 0A3 box. To our knowledge, we have developed the first mathematical model that incorporates all these regulatory components, accurately predicts the gene expression dynamics of *spo0A*, and estimates the regulatory roles, binding affinities, and interaction energies of the 0A boxes.

After testing different modes of promoter regulation, our best-fitting model predicts that P*v* and P*s* promoters are both purely kinetically regulated, i.e., the regulatory effect of 0A boxes do not influence the binding probability of RNAP, but rather modulate the subsequent transcription step to adjust the overall transcription rate (Figure 5A). In the case of the P*v* promoter, this can be explained by the 0A boxes being downstream the promoter region, which may cause the bound 0A∼P to act as a roadblock (44) to the progress of the transcription elongation complex, reducing the overall transcription rate (*v*_*max*_). A similar roadblock mechanism has been reported for the 0A∼P regulated *abrB* gene, where the 0A box region is also downstream of the promoter (45).

On the other hand, given that the 0A boxes are located upstream of the P*s* promoter, one would expect P*s* to be thermodynamically regulated. Surprisingly, the kinetic P*v* and thermodynamic P*s* regulation model fits worst the experimental data, with an optimization error twice as high as the best-fitting purely kinetic model (Supplementary Figure S4). However, supporting our results, in (46), single transcription assays revealed that the 0A boxes associated with *spoIIG* also have solely purely kinetic regulatory effects, despite being located upstream of the promoter region. Specifically, 0A∼P was shown to only stimulate the overall rate of transcription by affecting the post-closed-complex step and had no effects on the binding of RNAP to the DNA. To our knowledge, the exact mechanism responsible for the kinetic effects of upstream 0A boxes has not been identified. Namely, 0A∼P may act as a topological constrain for supercoils diffusion, inhibiting open complex formation, promoter escape, and transcriptional elongation steps (47). More experimental measurements are necessary to confirm this hypothesis.

Gene regulation by both kinetic and thermodynamic effects on a single promoter is also possible. In this case, occupancy of a TF binding site changes both the binding probability of RNAP and the maximum transcription rate. However, such model was not tested since a purely kinetic model is in accordance with the additivity property of the experimental data (Figure 3G).

Further, the existence of additivity in our experimental data is consistent with other experimentally validated models (48), which show that two tandem promoters express at a level between that of a single promoter and the sum of two single promoters. Lower expression than the sum of two single promoters is due to RNAP interference caused by the RNAP occupancy of the downstream promoter (48). As such, the perfect additivity of the experimental data suggests that both closed- and open-complex formation occur rather quickly in the P*s* promoter, preventing the interference between an elongating RNAP from P*v* and an initiating RNAP in P*s*.

In addition, the model predictions suggest that four 0A∼P molecules bind to each 0A box, although literature suggests 0A∼P binds as a dimer (38–40). One possible explanation for this discrepancy, is that 0A∼P may act similarly to LacI (49), i.e., functioning as a tetramer composed of two dimers. In this arrangement, one dimer binds to one 0A box, while the other dimer in the tetramer binds to a separate 0A box, possibly inducing DNA loop formation. More evidence is necessary to verify the tetrameric structure of 0A∼P transcription factor and the exact biomolecular events of the binding at 0A box(es).

We also note that the model predicted high-order interaction energies between 0A boxes differs from the *in vitro* EMSA estimations. While the fraction of bound DNA fragments in the EMSA data can be explained without interaction energy between 0A boxes, the biophysical model can only fit well assuming the existence of 0A box interaction parameters that are also key to understanding the saturation of 0A∼P binding. This may be explained by different DNA topologies in the two experiments without (for EMSA measurements) or with (for P*spo0A-lacZ* measurements) the presence of the *spo0A* promoter(s). The incorporation of the promoter(s) perhaps induces a change in the local chromatin state due to supercoiling accumulation (50), resulting in additional DNA interactions between bound 0A∼P.

Meanwhile, the model predicts that, as a group, the presence of 0A1-3 boxes has repressive effects on P*v* and P*s* promoters. However, at the individual level, 0A1 represses P*v*, 0A2 represses P*s*, and 0A3 is an activator of P*s* (Figure 6D). This is in agreement with past studies (23) suggesting that, at the onset of starvation, 0A3 is responsible for activating P*s* whereas 0A1 for repressing P*v*, flipping the switch from the vegetative to the sporulation promoter. Noteworthy, we did not observe a direct switching behavior, as suggested in (19), between the ‘P*s* strain’ and ‘P*v* strains’, i.e., we did not observe a simultaneous decrease and increase in ‘P*v*’ and ‘P*s* strain’ expression, respectively (Figure 2E and 2F, black lines). Rather, the activity of both promoters generally increased since the time of measurement, even with mutation of 0A boxes (Figure 2D-I), due to the strong activation caused by both σ^A^ and σ^H^-RNAP (Figure 7A-C). Nevertheless, ‘P*s* strains’ did show a higher fold increase over time (∼10.8 fold) when compared to ‘P*v* strains’ (∼1.4 fold).

Moreover, the regulatory effects of individual 0A boxes are in agreement with the results reported in (31). Interestingly, even though our model concludes that the effect of 0A∼P binding is relatively weaker than the increase in σ^A^ and σ^H^-RNAP to upregulate *spo0A* expression, 0A boxes were found to be necessary to induce proper sporulation (31). For the detailed analyses on how 0A box and promoter perturbations affect the phenotypes (biofilm formation and sporulation), we refer the readers to the separate work (31).

Finally, in this study, we do not directly investigate the role of 0A4 box on *spo0A* regulation. This is because 0A4 overlaps the -10 region of the P*s* promoter (11, 23). Consequently, it is not possible to mutate 0A4 box without disrupting the promoter sequence. As such, the reported effects of 0A1-3 boxes on P*s* (Figure 6D) may also be influenced by the presence of the 0A4 box. Nevertheless, past studies (23) have concluded that such effects are close to negligible. They found that, when P*s* promoter is substituted with another σ^H^-specific promoter (P*spoVG)*, the expressions of P*s* and P*spoVG* are similar with and without mutations to 0A2, 0A3, or the combination thereof (23), suggesting the presence of 0A4 box has no regulatory effect on P*s*.

Overall, our study introduces a new model that helps tackle one major challenge: the difficulty of predicting the regulatory outcomes of different regulatory configurations. Besides capturing the system’s phenomenological information (i.e., the observed input-output relationship between transcription factor concentration and gene expression), the model directly connects the experimentally adjustable parameters of the system (e.g. independent tuning of 0A boxes binding affinity and promoter affinity to RNAP) to its input-output response. As such, this model can be used to predict the dynamics of synthetically engineered strains that aim to fine tune the expression dynamics of the sporulation master regulator. Moreover, the model can also be applicable to study the regulatory role of other 0A boxes such as the ones regulating *spoIIA, spo0F*, the *sin* regulon, and *kinC* (12, 38, 51–54), as these genes encode proteins that play crucial roles regulating the biofilm production and sporulation.

## 4. Materials and Methods

### 4.1. Bacterial strains

The list of all strains used in this study is shown in Supplementary Table S1. All data from genetically perturbed *spo0A* strains were obtained from (31). In addition, we engineered two strains from *B. subtilis* DK1042 (NCIB 3610 background, BGSC3A38 as the *Bacillus* Genetic Stock Center strain number) (55). In one strain, we fused *lacZ* reporter to the WT P*citG* promoter. In the other strain, we fused *lacZ* reporter to the WT P*hyper-spac*. P*hyper-spac* used in this study is a modified form of the original P*hyper-spac* promoter that lacks the operator for the LacI repressor and is therefore constitutively active (9, 42). Each of the P*hyper-spac-lacZ* and P*citG-lacZ* (43) constructs was introduced into the competent biofilm-forming *B. subtilis* strain (DK1042, competent NCIB 3610) (55). The plasmids were constructed by using a DNA fragment containing the promoter region of *citG/hyper-spac* amplified with primers om454 and om455 using PY79 as the template. The resulting PCR fragment was digested with HindIII and BamHI and inserted into HindIII and BamHI digested pDG1728 (56).

### 4.2 Growth conditions

Briefly, lysogeny broth (LB) was used for growth of *B. subtilis*. 1.5% agar was used to make solid media plates. Cells harboring *lacZ* reporter fusions were cultured in liquid Minimal Salts glycerol glutamate (MSgg) medium with shaking at 37°C (26, 57, 58). Samples were collected hourly and assayed for β-galactosidase activities.

### 4.3 *β*-Galactosidase assay

Promoter activity of P*citG* and P*hyper-spac* was measured through β-Galactosidase assays performed as described in (59). Measurements were taken hourly from t = 2h until t = 9h. The first time point (t = 2h) corresponds to the onset of starvation. Promoter activity data of perturbed *spo0A* strains was obtained from (31) for the same time points.

### 4.4 Biophysical models

The framework of the biophysical models developed is based on previous models (32–34, 36, 37) where gene expression is considered to be dependent on the RNAP binding probability to the promoter. Adaptations of this model framework to predict the promoter activity levels of the single- (‘P*v* strains’ and ‘P*s* strains’) and dual-promoter mutant strains (‘P*s*P*v* strains’) are described in Supplementary Section II and III, respectively.

The statistical mechanical framework is applied to developed four models with different types of transcription control. The models are: 1) both P*v* and P*s* promoter are kinetically regulated; 2) both P*v* and P*s* are thermodynamically regulated; 3) P*v* is kinetically regulated, and P*s* is thermodynamically regulated; 4) P*v* is thermodynamically regulated, and P*s* is kinetically regulated.

In Supplementary Section IV, Subsection iv.1, we describe how the framework is applied to assume thermodynamic control of promoter activity by TFs. We first introduce the case for a single promoter present (Supplementary Section IV, Subsection iv.1.a) and then expand it for the dual-promoter system (Supplementary Section IV, Subsection iv.1.b).

Likewise, the adaptation of the statistical mechanical framework to assume kinetical control of promoter activity by TFs, for the single promoter system, is described in (Supplementary Section IV, Subsection iv.2.a). The expansion to the dual-promoter system is described in (Supplementary Section IV, Subsection iv.2.b).

Finaly, the description, for the two models in which there is mixed control (i.e. one promoter is thermodynamically regulated while the other is affected kinetically) is in Supplementary section IV, subsection iv.3. We note that all the derivations shown in Supplementary Sections II-IV assume the most complex case, in which three 0A boxes are present in the DNA.

Overall, for each of the models tested, the parameters of *Eqn. 4* reflect the different types of regulatory control the 0A box has on the promoter and the biophysical mechanism of action (repression or activation) on each promoter. For purely thermodynamic regulation, the effective transcription rate comes from the effect bound 0A∼P has on the binding probability of RNAP to the promoter. Whether the 0A box is bound or unbound has no effect on the maximum transcription initiation rate. Conversely, for a purely kinetically controlled promoter, 0A∼P binding influences the post-closed-complex step, therefore modulating the maximum transcription initiation rate.

### 4.5. Model fitting procedure

We started by constraining the model parameters by using previous literature results or estimations from experimental data. Namely, we assumed wild-type 0A∼P dynamics (i.e., [0*A*∼*P*] in Supplementary *Eqn. II.4*) to follow the same dynamics as in (26). Dynamics shown in Supplementary Figure S3. In addition, the binding energy of 0A boxes was estimated from EMSA data and reconciled with *in vivo* measurement with an additional unknown parameter (Supplementary section V). The cooperativity estimated from EMSA (*N*) is directly used in the biophysical model (*N*_0 *A*_ in Supplementary *Eqn. II. 4* and *Eqn. III. 9-11*).

All other variables in the model, including the interaction between RNAP and 0A∼P, the maximum transcription initiation rate of different configurations, as well as the concentration of RNAP recognized by P*v* and P*s*, are unknown and predicted from fitting the models to the *spo0A-lacZ* promoter activity measurements over time (Figure 2D-I). For example, to explain all experimental data of a mutant strain with one 0A box (such as 0A1), we fit *Eqn. 4* describing P*v* promoter to the ‘P*v* strain, 1’ experimental data, *Eqn. 4* describing *Ps* promoter to the ‘P*s* strain, 1’ data, and Supplementary *Eqn. III.18* to the ‘P*v*P*s* strain, 1’ data for every possible concentration of 0A∼P.

Each of the four models was simulated under the same number of parameters (see Supplementary Table S3 for a full list of parameters) for at least 20 independent iterations.

To find the best fitting model to the experimental data, we minimize the difference between model computed data and experimental data (*E*) using the following objective function:

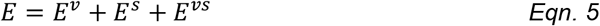

Where,

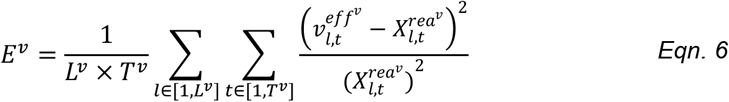

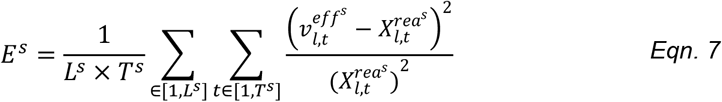

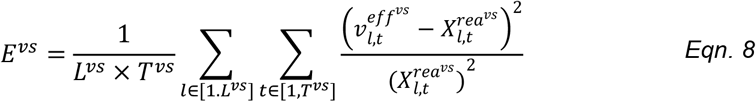

Where *L*^*i*^ is the total number of mutant strains, *T*^*i*^ is the number of hours measured, and 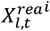 is the mean experimental (“real”) promoter activity of a particular mutant strain (*l*) at a particular time point/0A∼P concentration (*t*) across several biological replicates. The superscripts *i* = {*v*; *s*; *vs*} represent the types of promoter(s) being present. There are eight mutant strains in each cohort (*L*^*v*^ = *L*^*s*^ = *L*^*vs*^ = 8) and every strain is measured for eight hours (*T*^*v*^ = *T*^*s*^ = *T*^*vs*^ = 8). MATLAB (R2021a) Particle Swarm Optimization algorithm was used to minimize the error (60).

## 5. Data availability statement

A data software package with the statistical thermodynamic model used in Results section 2.2, the four biophysical models tested, model fitting scripts (Results section 2.3), and figure/table generation scripts are deposited in GitHub (https://github.com/yjzhang3/Biophysical-model-spo0A-transcription-regulation.git). The experimental data

## 6. Supplemental data

Supplemental data available online.

## 7. Author contributions

O.I and M.F conceived and supervised the study. Y.Z. executed the experimental data analysis, and model design implementation and simulation, to which C.S.D.P contributed to. Z.C designed and implemented the model to determine 0A∼P dynamics. B.Z and M.F planned and executed the experimental measurements. Y.Z. and C.S.D.P. drafted all documents which were revised by all co-authors.

## 8. Acknowledgements

This work was supported by the National Science Foundation (MCB-2204402 to OAI (co-PI), PI: Jeff Tabor), the Welch Foundation (Grant C-1995 to OAI and E-2099 to MF), and by the Jenny and Antti Wihuri Foundation (to C.S.D.P.).

## SUPPLEMENTAL INFORMATION FOR

### (I) Model fraction of DNA bound by 0A∼P

In this section, we describe the data processing of the electrophoresis mobility shift assay (EMSA) used to examine the properties of 0A boxes and constrain the biophysical model.

#### (i.1) Computing fraction of bound DNA using law of mass action under equilibrium

The process of a particular 0A box being bound by 0A∼P can be described according to the reaction scheme *Eqn. I. 1*, with the forward reaction rate *k*_1_ and backward reaction *k*_2_:

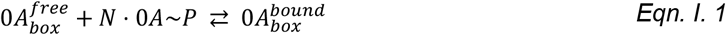

Here, as in the main manuscript, 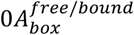 stands for free/bound binding site conformation, respectively, and *N* is the number of free 0A∼P molecules that bind to the free site. In accordance with the mass-action equilibrium law one can write the dissociation constant *K*_*d*_ as:

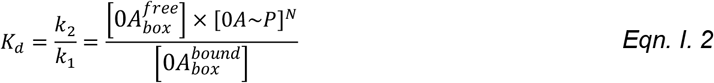

Assuming the total concentration of the binding site to be 1, the sum of bound and unbound form can be written as:

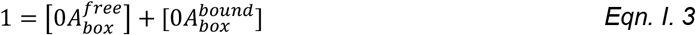

Then, by combining *Eqn. I. 2* and *Eqn. I. 3*, the concentration of bound binding sites can be written as:

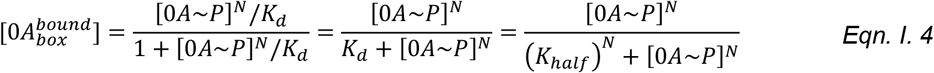

We chose to report binding affinity in the same unit of the 0A∼P concentration. Therefore, we let *K*_*d*_ = (*K*_*half*_)^*N*^, where *K*_*half*_ is the 0A∼P concentration resulting in half-maximal level (i.e., probability that a 0A box is bound is 50%).

### (i.2) Augmenting EMSA data

To quantify the fraction of DNA bound to 0A∼P, we performed gel electrophoresis mobility shift assay (EMSA) for DNA fragments with mutagenesis combinations of the 0A boxes (1). These fragments include those with single boxes (0A1, 0A2, and 0A3) and those with multiple boxes (0A12, 0A13, 0A23, 0A123). For each DNA fragment, the experiment was conducted over 11 concentrations of 0A∼P (0 to 2 *µM*, at an interval of 0.2) with *M* biological replicates, with *M* ∈ {2,3,4}.

To model the fraction of bound DNA 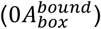 for strains with only one 0A box (“1”, “2”, “3”, denoting the presence of 0A boxes), we performed the augmentation protocol for these three DNA fragments. The protocol is as follows:

1. From the biological replicate data, we compute the mean and standard deviation of the fraction of bound DNA, at each 0A∼P concentration. We then randomly sampled from a Gaussian distribution with the same mean and standard deviation to obtain the augmented data point at each corresponding 0A∼P concentrations.
2. Given that, for each DNA fragment, there are a total of 11 × *M* experimental data points before augmentation. Using the random sampling method above, we also sampled 11 × *M* data points, for each DNA fragment.
3. Combining augmented datapoints for all three types of DNA fragment, this leads to 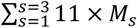 newly generated data points, where *M*_*s*_ is the number of replicate experiments for each type of DNA fragment “*s*". We consider these many data points as one augmented dataset.
4. We repeated the protocol 1000 times, leading to 1000 augmented EMSA datasets in total.

#### (i.3) Fitting the mathematical expression to EMSA data

Given known concentrations of 0A∼P, we want to solve for half saturation constant (*K*_*half*_) and the number of 0A∼P molecules (*N*). To this end, we fit *Eqn. I. 4* to a single augmented dataset of the single-0A box measurements. The optimization error between model prediction and augmented empirical EMSA data (*E*_*EMSA*_) is defined as:

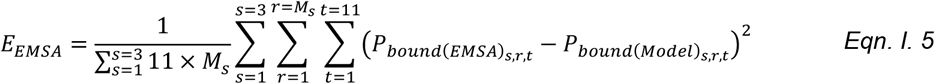

Where, *s* is the type of DNA fragment (“1”, “2”, “3”), *r* is the particular replicate out of *M*_*s*_ total replicates for each DNA fragment (*M*_*s*_ ∈ {2,3,4} depending on *s*), and *t* is the concentration of 0A∼P unique to each measurement. 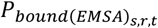 and 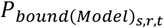 is the experimental data and model-generated data, at a specific *s, r, t*, respectively.

A total of 1000 sets of *K*_*d*_ parameters, including *K*_*half*_ for each 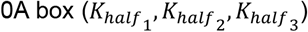 are obtained by fitting *Eqn. I. 4* to all 1000 augmented datasets, one at a time. Model fitted to each augmented dataset is optimized at least 20 times using MATLAB Particle Swarm Optimization algorithm (2). The mean with 95% confidence interval in each optimized *K*_*half*_ parameter across 1000 augmented sets is shown in Figure 4B. We also examined the model fits by examining different possible values of *N*, and only *N* = 4 fits the best (Table S2).

#### (i.4) Developing a general statistical thermodynamic model and predicting fraction of bound DNA for strains with multiple binding sites

In the previous section i.1, we showed that the concentration of bound DNA can be computed using the law of mass action under equilibrium, assuming the total concentration of DNA is 1. This approach is equivalent to computing the probability of DNA being bound (*P*_*bound*_) using equilibrium statistical thermodynamics.

Specifically, to compute *P*_*bound*_, we divide the statistical weight representing 0A∼P bound by the sum of the statistical weights representing all possible system states (3, 4). For a DNA fragment with only one 0A box present, only two possible system states are possible: 0A∼P bound to the 0A box, or 0A∼P free. As a result, the probability of a DNA fragment with one 0A box being bound by 0A∼P *(P*_*bound, one box*_) is given by *Eqn. I. 6*, where the statistical weight of each state depends on the binding energy at a particular 0A box (*G*_*i*_) and the cooperativity (*N*) (5, 6). The unbound state is assumed to have zero binding energy.

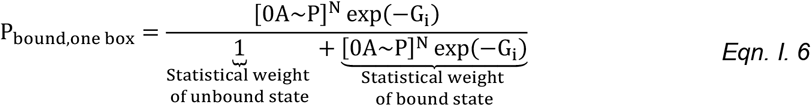

Noteworthy, comparing *Eqn. I. 4* and *Eqn. I. 6*, we see that, for a 0A box ‘*i’:*

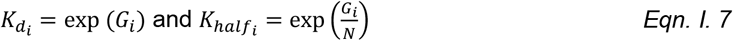

Where *i* ∈ [1,2,3]

To understand why this is true intuitively, consider the case where, in *Eqn. I. 6*, a more negative binding energy increases the statistical weight of the bound state to increase the probability of DNA being bound. When the binding energy is more negative, *K*_*d*_ become smaller, suggesting the forward reaction (DNA is bound) is favored and the binding affinity is higher.

Next, we derive statistical thermodynamic expressions to compute the probability of DNA being bound by 0A∼P for DNA fragments with multiple 0A boxes. For DNA fragments with two 0A boxes, box *i* and *j*, there are four possible states the DNA is bound by 0A∼P: 1) neither *i* or *j* bound; 2) *i* bound; 3) *j* bound; 3) *i* and *j* bound. As such the cumulative probability where at least one 0A box is bound (*P*_*bound,two boxes*_) can be written as:

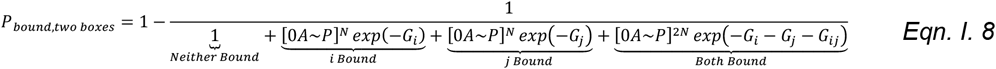

Where, *G*_i_ and *G*_*j*_ are the binding energies of the each 0A box, and *G*_*ij*_ is the interaction energy between two bound transcriptional factors (also known as “secondary interaction energy”). The variable *N* represents the cooperativity level between 0A∼P molecules. The underlying brackets indicate the type of configuration represented by the Boltzmann weight.

For DNA fragments with three 0A boxes, box *i, j*, and *k*, there are 2^3^ possible configurations the DNA fragment is bound by 0A∼P. Similarly, one can calculate the cumulative probability where at least one 0A box is bound (*P*_*bound,three boxes*_) as:

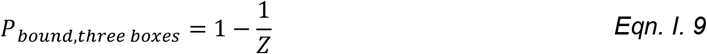

Where, *Z* = *Z*_1_ + *Z*_2_ + *Z*_3_ + *Z*_4_. The decomposition of *Z*_1_, *Z*_2_, *Z*_3_ and *Z*_4_ is shown below:

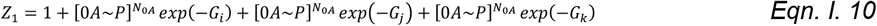

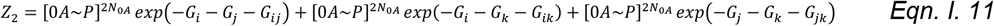

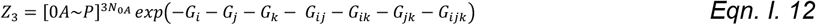

Noteworthy, in *Eqn. I. 9*, in addition to secondary energy between every pair of 0A boxes (*G*_*ij*_), the tertiary interaction energy (*G*_*ijk*_) when all three 0A boxes are bound is also considered. Particularly, *Z*_1_ is the sum of statistical weight for configurations with only one 0A box occupied; *Z*_2_ represent all statistical weights for configurations with two 0A boxes occupied. *Z*_3_ is the statistical weight of the configuration with all three boxes being occupied.

We then use each set of model-optimized *K*_*d*_ parameters from section i.3 to compute the probability of DNA being bound, for strains with two or more 0A boxes (“12”, “13”, “23”, and “123”). To start with, we convert *K*_*d*_ parameters to binding energy *G* based on *Eqn. I. 7*. Then, assuming no secondary or tertiary energy, i.e., *G*_*ij*_ = 0 and *G*_*ijk*_ = 0, we compute the probability of DNA bound using *Eqn. I. 8* and *Eqn. I. 9*. The predicted results are shown in Supplementary Figure S2.

### (II) Biophysically model promoter activity for DNA with a single promoter

The single-promoter model framework is applied to strains with only P*v* present (“P*v* strains”), and only P*s* present (“P*s* strains”).

For a DNA fragment with a single promoter and with *n* − 1 ligand sites, binding of transcription factors (TFs) and transcriptional machinery results in 2^n^ possible unique ways in which these *n* sites can be occupied. Each possible way is defined as a “configuration” (*α*), which is represented using binary variables (σ):

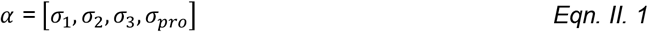

Where *σ*_*i*_ = 1 denotes the site being occupied, and 0 otherwise. In this specific model, the variables *σ*_1_, *σ*_2_, *σ*_3_ represents bound/unbound 0A1-3 box, respectively. The variable *σ*_*pro*_ represents bound/unbound promoter by RNAP.

Next, for the configurations that can transcribe i.e., those with *σ*_*pro*_ = 1, the effective transcription rate (*v*^*effective*^) is modeled as the product of RNAP binding probability (*P*) and the maximum transcription initiation rate (*v*_*max*_) (6–9). The effective transcription rate of each mutant strain (*MS*) considers the contributions from all promoter-bound configurations, and is therefore calculated as:

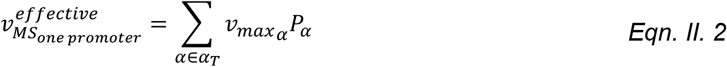

Where, 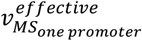, is the effective transcription rate of a mutant strain with a single promoter. Variable, *α*_*T*_, is the number of possible configurations with promoter bound, i.e., *α*_*T*_ *=* [*σ*_1_, *σ*_2_, *σ*_3_, *σ*_*pro*_ = 1].

Assuming both reactions happen at thermodynamic equilibrium, in the below sections we expand *P*_*α*_ and *vmax*_α_ using dimensionless energy landscape parameters (Figure S11) (5, 6).

#### (ii.1) Binding probability, *P*_*α*_

The binding probably of RNAP to the promoter can be written as the ratio between the Boltzmann weight (*BW*) of a particular configuration (*α*_*T*_) and the sum of Boltzmann weights over all configurations (also known as the partition function) (5–7, 10), as shown in *Eqn.II.3*:

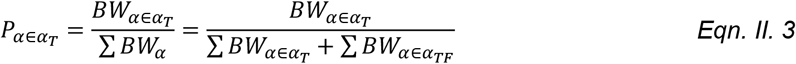

Here, *α*_*TF*_ corresponds to the configurations where RNAP is not bound to the promoter (i.e., *α*_*TF*_ = [*σ*_1_, *σ*_2_, *σ*_3_, *σ*_*pro*_ = 0]). The Boltzmann weight of each configuration (*BW*_*α*_) depends on the free energy of the configuration, the concentration of the bound ligands, and the cooperativity of the ligands. As such, one can further expand *Eqn. II.3* as:

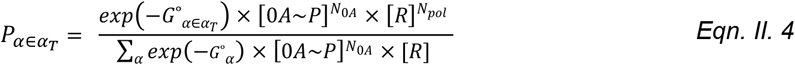

Where, [*R*] and [0*A*∼*P*] are the concentration of RNAP and Spo0A∼P, respectively. Variable *G*°_*α*_ is the free energy of configuration *α*. The variables *N*_0*A*_ and *N*_*pol*_ are the cooperativity of bound Spo0A∼P molecules and cooperativity of bound RNAP, respectively. We assume *N*_*pol*_ = 1, given that not more than one RNAP can bind simultaneously to a single promoter region. *N*_0*A*_ is assumed to be 4 based on the estimation of EMSA data (Result Section 2.2; Supplementary Section I).

Further, the denominator of *Eqn. II.4*, can be decomposed into configurations with promoter bound (*α*_*T*_) and promoter unbound by RNAP (*α*_*TF*_) :

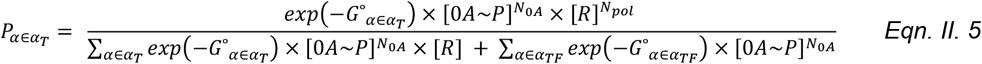

The free energy of a configuration, *G*°_*α*_, depends on the binding of ligand sites and can be written as:

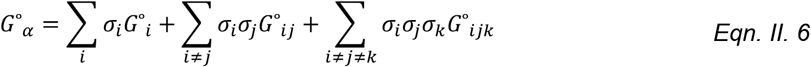

With, *i, j* and *k ϵ* {1,2,3, *pro*}.

Here, *G*°_*i*_, *G*°_*ij*_, and *G*°_*ijk*_ are the binding energy of each bound ligand, secondary interaction energies between each pair of bound ligands and tertiary interaction energies between every three of bound ligands, respectively.

According to *Eqn.II.6*, each single-site binding energy can only contribute to *G*°_*α*_ if the corresponding site is occupied (*σ* is equal to 1). Moreover, each interaction energy can only contribute to *G*°_*α*_ if the cognate binding sites are all occupied (product of *σ*’s is equal to 1). Notably, we assume the free energy for the configuration with no site being bound ([*σ*_1_ = 0, *σ*_2_ = 0, *σ*_3_ = 0, *σ*_*pro*_ = 0]) to be 0.

To make notations comprehensive for the following sections, below we show how to expand the summation terms in *Eqn. II. 6* as a function of specific energy parameters:

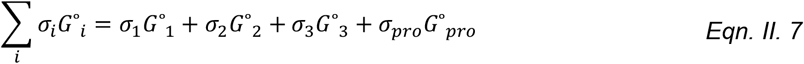

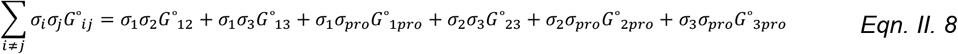

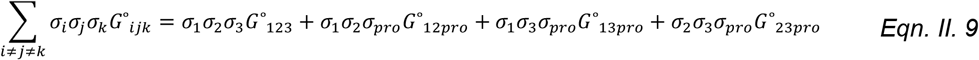

Where *G*°_1_, *G*°_2_, *G*°_3_, and *G*°_*pro*_ are binding energies of each ligand binding site; *G*°_12_, *G*°_13_, *G*°_23_, and *G*°_123_ are interaction energies between 0A∼P bound at each 0A box; *G*°_1*pro*_, *G*°_2*pro*_, *G*°_3*pro*_, *G*°_12*pro*_, *G*°_13*pro*_, and *G*°_23*pro*_ are interaction energies between at least one 0A∼P and the RNAP bound at the promoter.

To emphasize, for those configurations where promoter is bound (*α* ∈ *α*_*T*_), the free energy of these configurations 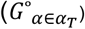 assumes *σ*_*pro*_ = 1. Conversely, the free energy of all promoter-unbound configurations (*G*°_*α*∈*TF*_) assumes *σ*_*pro*_ = 0.

#### (ii.2) Maximum initiation rate, *v*_*max*_

The maximum transcription rate of configuration *α* can be written as:

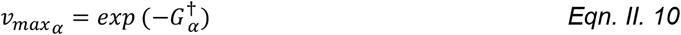

Where, 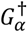 is the activation energy for the transcription reaction. 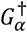 can be decomposed as:

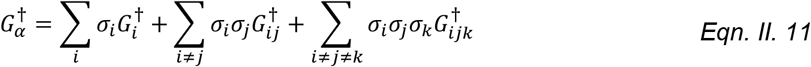

With, *i, j* and *k ϵ* {1,2,3, *pro*}.

Here, 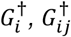, and 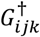 are the activation energy resulted from each site being bound, activation energy resulted from a pair of sites being bound, and activation energy from three sites being bound, respectively. Further, we assumed that 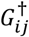 and 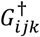 are nonzero.

According to *Eqn. II. 11*, each single-site binding energy can only contribute to 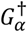 if the corresponding site is occupied (*σ* is equal to 1). Also, each interaction energy can only contribute to 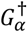if the cognate binding sites are all occupied (product of *σ’*s is equal to 1).

### (III) Biophysically model promoter activity for DNA with two promoters

The model framework described above (Supplementary Section II) can be extended to compute gene expression for strains with both P*v* and P*s* promoters present (“P*v*P*s* strain”). For a gene segment with two promoters and three transcription factor binding sties, each possible configuration (α) can be represented as:

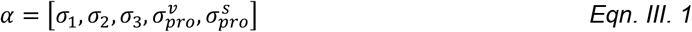

Where, 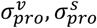 denote P*v* bound/unbound or P*s* bound/unbound, respectively.

In a two-promoter system, transcription is possible if at least one promoter is bound. Therefore, configurations where only P*v* is bound (*α*_*v*_), configurations where only *Ps* is bound (*α*_*s*_), and configurations where both P*v* and P*s* are bound (*α*_*vs*_) all contribute to the effective transcription rate. As such the effective transcription rate of a two-promoter mutant strain 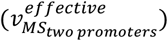can be written as:

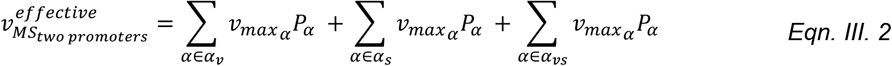

Where,

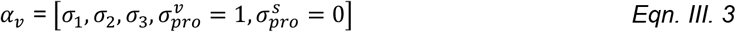

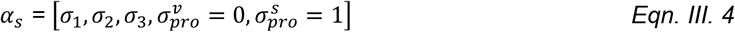

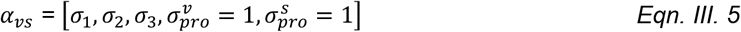

Similarly to *Eqn.II.3*, the probability of RNAP binding to a particular configuration, *P*_*α*_, is computed using Boltzmann weights of configurations (*BW*_*α*_) (5–7, 10). This way, one can explicitly express *P*_*α*_ depending on whether the configuration has P*v* bound (*α*_*v*_), P*s* bound (*α*_*s*_), or P*v and* P*s* both bound (*α*_*vs*_), as shown below:

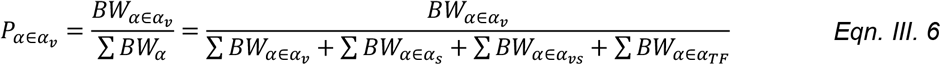

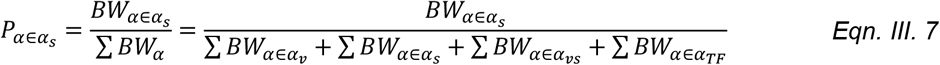

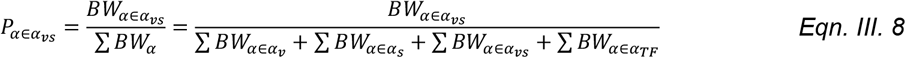

Since Boltzmann weights of each configuration depends on the free energy of the configuration, the concentration of the bound ligands, and the cooperativity of the ligands, one can further expand *Eqn.III.6-8* as:

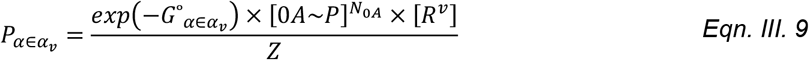

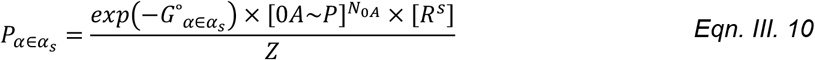

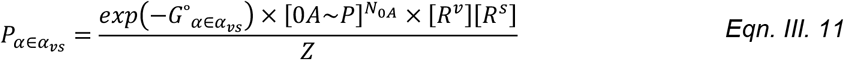

Where [*R*^*v*^] and [*R*^*s*^] are the concentration of RNAP bound at promoter *Pv* and *Ps*, respectively. Meanwhile, *Z*, the statistical weight of all configurations, can be written as:

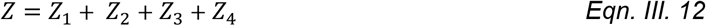

Where,

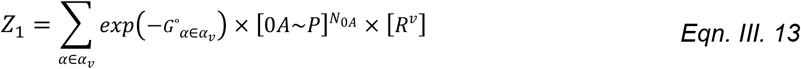

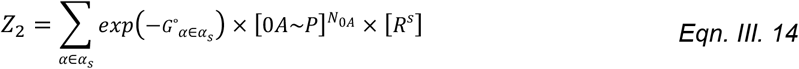

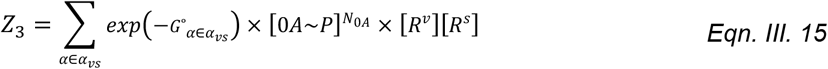

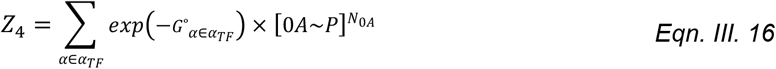

Similar to *Eqn. II. 6*, for a two-promoter system, *G*°_*α*_, depends on the binding of ligand sites and can be written as:

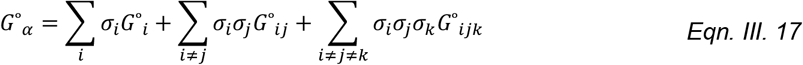

With, *i, j* and *k ϵ* {1,2,3, *pro*^*v*^, *pro*^*s*^}

The summation terms here can be further decomposed in specific energy parameters as in *Eqn. 7-9*. However, due to the existence of 2^5^ (three 0A boxes and two promoters) total possible configurations, the decomposition becomes too extensive to be shown.

Also, common existing parameters between *Eqn. III.17* and *Eqn. II.6* have the same value (e.g., the interaction energy of 0A1 and 0A2 box, *G*°_12_, is assumed to be the same, independently of how many promoters the mutant strain has). Noteworthy, the models do not assume the existence of interaction energy between the two promoters, with or without 0A boxes bound.

Lastly, due to the additivity observed in the empirical data, we assume that, for configurations with the same 0A boxes being occupied, *v*_*maxα*_ is additive between the two promoters. As such, *Eqn. 2* can be rewritten as:

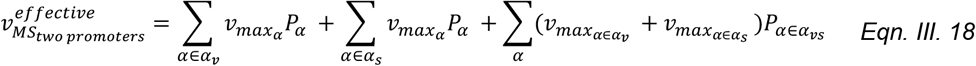

Where, 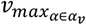 and 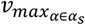 are the maximum transcription initiation rates unique to *Pv* and *Ps* promoter, respectively. The values of 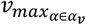 and 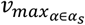 are the same as 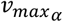 for each respective single promoter model. Noteworthy, this assumption does not necessarily result in an additivity level equal to 1 for all the biophysical models tested.

We also note that in main manuscript, for simplicity, Section 2.5 configurations are shown as a 3-digit number. These configurations all assume promoter(s) are bound. That is, notation ‘101’ for ‘P*v* strains’ in the main manuscript is in fact 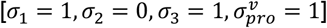. Similarly, notation ‘100’ for ‘P*v*P*s* strains’ in the main manuscript is in fact 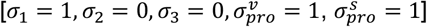.

### (IV) Biophysical models with different types of transcription control

In the previous section (Supplementary section II and III), we formulated the biophysical framework to model the effective transcription rate of a gene regulated by a single- or dual-promoter system. We assume TF can affect promoter activity by two different control mechanisms: thermodynamic or kinetic (Supplementary Figure S12 and S13). In the following section, we introduce how each mechanism is quantitively represented. Then, within each type of control, we further describe how it is applied to a single- and dual-promoter system. Noteworthy, since 0A∼P is the transcription factor in this case, we use “TF” and “0A∼P” interchangeably, and “TF binding site” and “0A box” interchangeably.

#### (iv.1) Biophysical model with purely thermodynamic control

A purely thermodynamic model assumes TF binding affects the probability of RNAP binding to the promoter but has no impact on the overall transcription rate (Supplementary Figure S12). Below we show how we applied such assumption to the single- and dual-promoter system.

##### (iv.1.a) For a single-promoter mutant strain

In our study, the single promoter system corresponds to single-promoter mutant strains with either P*v* or P*s* promoter. Since thermodynamic control has no effect on the overall transcription rate, the 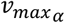, in *Eqn. II.2*, has the same value for all configurations (*α*).

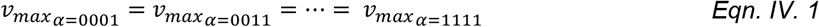

Instead, TF binding changes the transcription rate by affecting the probability of RNAP binding, specifically, by assuming nonzero interaction energy between any bound 0A∼P and bound RNAP. These include 0A∼P/RNAP secondary energies (*G*°_*ij*_, *i* ∈ {1; 2; 3}, *j* ∈ {*promoter*}) and 0A∼P/0A∼P/RNAP tertiary energies (*G*°_*ijk*_, *i* ∈ {1; 2; 3}, *i* ≠ *j* ∈ {1; 2; 3}, *k* ∈ {*promoter*}). The sign of these interaction energies determines the mechanism of action as described below:

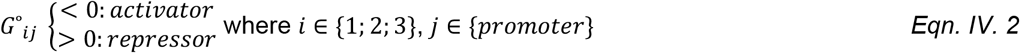

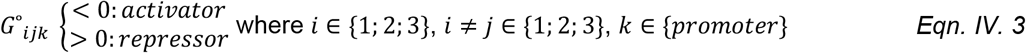

Given that all maximum transcription rates are identical regardless of the configuration, *Eqn. II. 2* for “P*v strains*” and “P*s strains*” can be simplified into:

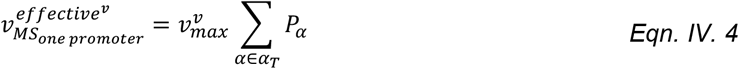

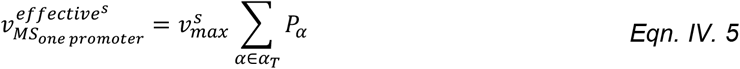

Where 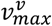 and 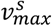 are maximum overall transcription rates for “*Pv strains*” and “*Ps strains*”, respectively.

##### (iv.1.b) For a double-promoter mutant strain

In our study, the double-promoter system corresponds to the mutant strains with both *Pv* and *Ps* promoters, ‘P*v*P*s strains*’. Given *Eqn. III. 18* and *Eqn. IV.4-5*, the two-promoter effective transcription rate of a mutant strain with two promoters, under purely thermodynamic control, can be described as:

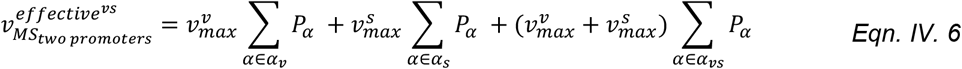

#### (iv.2) Biophysical model with purely kinetic control

A purely kinetic model assumes TF binding affects the overall transcription rate but has no impact on the recruitment of RNAP to the promoter (Supplementary Figure S13). Below we show how we applied such assumption to the single- and dual-promoter system model framework.

##### (iv.2.a) For a single-promoter mutant strain

In the kinetic control mechanism, the RNAP binding probability is not affected by 0A∼P binding. As such, there is no interaction between RNAP and 0A∼P. Instead, for kinetic control, the effect of TF binding is on the maximum transcription rate. As such, 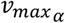 in *Eqn. II. 2* is assumed to be different for 0A∼P bound and 0A∼P unbound configurations.

In addition, the relative magnitude of 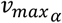 reflects the mechanism of action. If 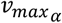 of a 0A∼P bound configuration is smaller than the 0A∼P unbound configuration, the binding of the 0A box has a repressive effect, or vice versa, as described below:

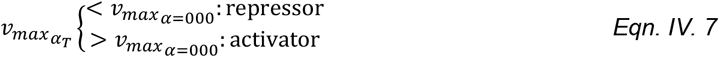

Where, *α*_*T*_ = [*σ*_1_, *σ*_2_, *σ*_3_, *σ*_*promoter*_ = 1].

The decomposition of the effective transcription rate *(Eqn. II.2*), for the kinetic control model, is described in the following paragraphs.

First, given that secondary (*G*°_*ij*_) and tertiary interaction (*G*°_*ijk*_) energies between bound 0A∼P and bound RNAP are null (zero) in a kinetic model, *Eqn. II.6* can be written as follows for promoter-bound (*α*_*T*_, where *σ*_*pro*_ = 1) and promoter-unbound (*α*_*TF*_, where *σ*_*pro*_ = 0) configurations, in accordance with *Eqn. II. 7-9*:

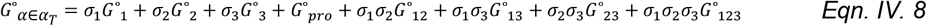

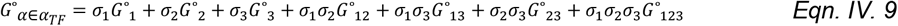

Note that *Eqn. IV.8* can also be written as a function of *Eqn. IV.9*:

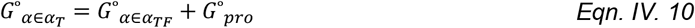

As such, the binding probability of RNAP to the promoter, *Eqn. II.5*, becomes:

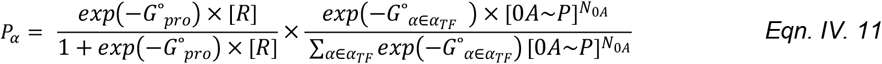

Note that the factorization of the term 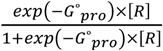 suggests the effect of RNAP is independent from the effect of transcription factor on binding probability, as expected in a purely kinetic model.

Finally, the effective transcription rate of a mutant strain 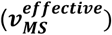, *Eqn. II.2*, can be written as:

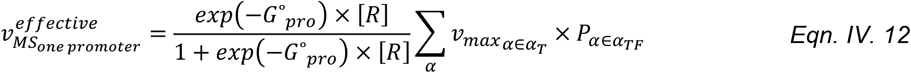

Where, the rate can be computed by summing the binding probability of 0A∼P (*α* ∈ *α*_*TF*_), weighted by their corresponding 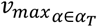 (for example, probability of the configuration [*σ*_1_ = 1, *σ*_2_ = 0, *σ*_3_ = 1] will be multiplied by the *v*_*max*_ of configuration 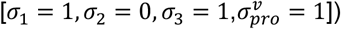), and multiplied by the factorization term.

##### (iv.2.b) For a double-promoter mutant strain

Below we show how to mathematically formulate the effective transcription rate of a dual-promoter system assuming TF effects are kinetic. We also show analytically that a purely kinetic control model leads to an additive transcription rate between two promoters.

First, in the dual-promoter system, the free energy for configurations with no promoter bound 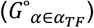 are equal to that in the single-promoter system (*Eqn. IV.9*). Further, since 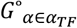 does not account with interactions between 0A∼P and RNAP bound to either promoter, and given there is no interaction between the two bound promoters, the energy for each group of configurations, 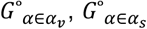, and 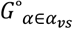, in *Eqn. III.9-11*, can be conveniently expressed as a function of 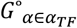.

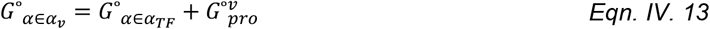

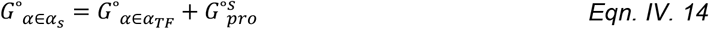

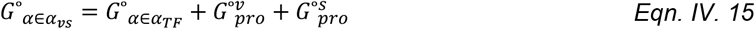

With this simplification, the binding probabilities in *Eqn. III. 9-11* can be written as:

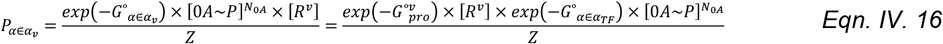

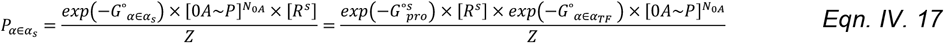

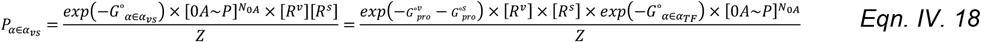

Given *Eqn. IV.13-15, Z*, the statistical weight of all configurations, can be written as:

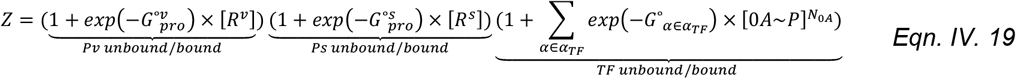

In accordance with *Eqn.III.2*, the effective transcription rate 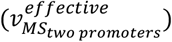 is then each promoter-specific configuration weighted by their promoter-specific 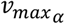:

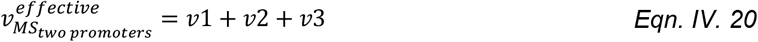

Where, *v*1, *v*2, and *v*3 can be written as:

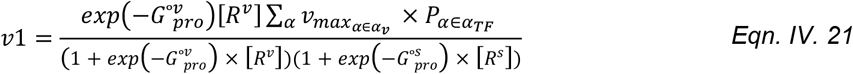

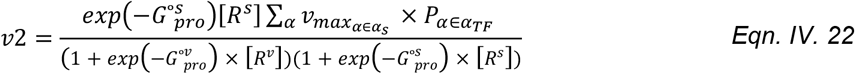

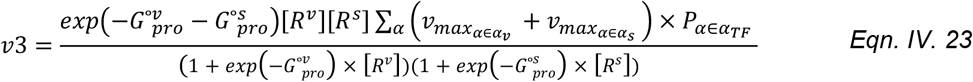

Here, 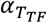 are the set of configurations concerning the binding of TF only. The variables 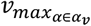 and 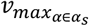 are the maximum transcription rate for each promoter-specific configuration when promoter is bound.

Further, one can expand the numerator of *v*3 (*Eqn. IV.23)* into:

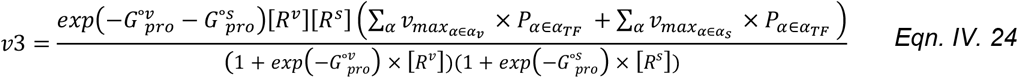

As such, now *Eqn. IV.20* be rearranged into

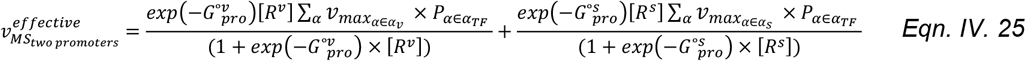

Which is exactly the sum of single-promoter effective transcription rate (*Eqn. IV.12*) between two promoters. This additivity property of the purely kinetic model is in agreement with the additivity observed in the experimental data.

#### (iv.3) Model with mixed control

The types of transcriptional control described in Supplementary Section iv.1 and iv.2 assume 0A∼P acts on both promoters with the same type of control: thermodynamically or kinetically. It is also possible that transcriptional regulation at each promoter is unique, including two possibilities: i) kinetic P*v* and thermodynamically regulated P*s*; ii) thermodynamic P*v* and kinetically regulated P*s*. Below we introduce the model framework of single- and double-promoter models with mixed control, assuming the case of kinetic P*v* and thermodynamically controlled P*s*.

For simplicity, the framework for single-promoter thermodynamic model and for single-promoter kinetic model have been developed in Supplementary Section iv.1.a and Supplementary Section iv.2.a (*Eqn. IV. 5* and *Eqn. IV.12*) and will not be detailed here.

To compute the effective transcription rate of a dual-promoter mixed model, we follow the general framework introduced in Supplementary Section III to calculate the RNAP binding probability. Using *Eqn. III.6* (generalized form of the Boltzmann factor for P*v* bound configurations), the contribution from P*v* promoter due to kinetic control can be written using the same rationale as in Supplementary Section iv.2.b:

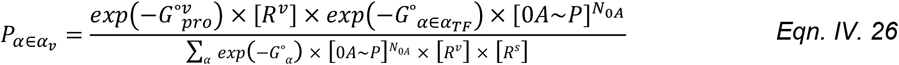

We then assign a maximum transcription initiation rate for all configurations in the two-promoter system. Based on Supplementary Section iv.1, because P*s* is thermodynamically regulated, all 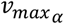 for *α* ∈ *α*_*s*_ are the same (here denominated as universal 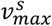). Based on Supplementary Section iv.2, the kinetically-regulated P*v* will have unique *v*_*maxα*_ for each configuration. In accordance with *Eqn. III. 18*, the effective transcription rate for a dual promoter mutant strain can be written as:

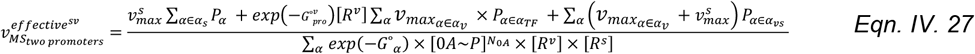

### (V) Constrain biophysical gene regulatory model with EMSA predicted parameters

In this section, we show how parameters predicted from EMSA data are used to constrain the biophysical gene regulatory model.

In Supplementary Section I, we obtained binding dissociation constant for each 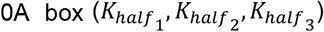 and *N*. We first converted optimized *K*_*half*_ parameters into binding energy parameters according to *Eqn. I. 7*, and call these 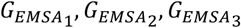. Then, we let the binding energy of single 0A boxes (*G*°_1_, *G*°_2_, *G*°_3_, see *Eqn. II. 7*) in the biophysical model equal to 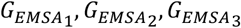, respectively. In other words, the occupancy of 0A boxes during transcription is assumed to be the same as the occupancy of 0A boxes during EMSA.

On the other hand, the parameter *N* is used as it is. We let cooperativity of 0A∼P (*N*_0*A*_ in *Eqn. II. 5*) equal to EMSA-predicted *N*.

However, directly using 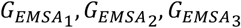 in the biophysical model fails to yield a good fit. We hypothesize this is because not all 0A∼P were phosphorylated during EMSA, i.e., the actual concentration that binds to 0A boxes are lower than the reported Spo0A concentration added to the DNA fragment (0∼2 µM) during gel shift experiment. Consequently, an overestimation of ligand concentration will underestimate the binding affinity of the substrate. To this end, we introduced an extra parameter, *α*, to reflect the percent of phosphorylation of Spo0A *in vitro*. This successfully reconciled the EMSA and promoter activity datasets, leading to a best-fit model. Below we show how this is done.

Let [0*A*∼*P*]_*in vitro*_ be the reported, overestimated 0A∼P concentration from EMSA, and [0*A*∼*P*]_*in vivo*_ be the 0A∼P concentration during transcription of *spo0A*. Assuming that *α* percent of [0A∼P]_*in vitro*_ during EMSA binds to 0A and reasonably describes the *in vivo* condition, one can write:

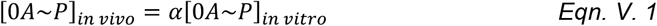

Next, let *G*_*in vitro*_ be the *in vitro* binding energy of a 0A box estimated from EMSA data, and *G*_*in vivo*_ be the *in vivo* binding energy (which is used the biophysical model). Based on *Eqn. I. 6*, the probability of DNA being bound by 0A∼P under *in vivo* conditions can be written as:

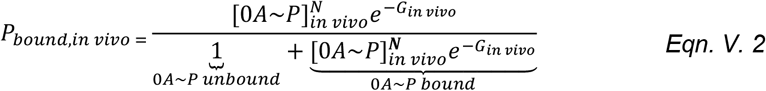

After correcting for the ligand concentration, during EMSA, the probability of DNA being bound by 0A∼P can be rewritten below using *Eqn. I. 6 and Eqn. V.1* as a function of *in vitro* 0A∼P concentration:

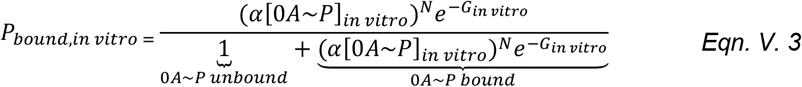

Assuming *Eqn. V.2* is equal to *Eqn. V.3*, that is, assuming the phosphorylation of *in vitro* 0A∼P concentration can correct for the *in vitro* binding energy:

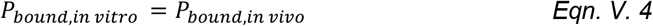

One can derive the quantitative relationship between *G*_*in vitro*_ and *G*_*in vivo*_ as:

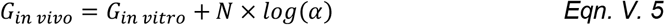

According to *Eqn. V. 5*, the *in vivo* binding energy relates to *in vitro* energy by a constant that is a function of *α*. Based on this, we can constrain the biophysical model using *in vitro* binding energies by adding an extra parameter that represents the constant *N* × log(α). We refer to this constant as *β*. Specifically, each binding energy in the biophysical model (*G*°_1_, *G*°_2_, *G*°_3_) can now be represented as 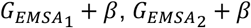, and 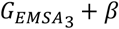. Optimizing three unknown binding energies in the biophysical model are now reduced to optimizing *β* only.

In the best-fit biophysical model, *α* was found to be 20%.

### (VI) Compute 0A box occupancy probability using best-fit kinetic model parameters

In Result Section 2.4, we showed that the effect of 0A∼P binding is saturated at t = 2 h. Here, we first demonstrate how to compute the probability of 0A∼P binding to 0A boxes. Then, we show why saturation in 0A box binding probability leads to no change in effective transcription rate as a function of 0A∼P concentrations.

The occupancy probability of 0A boxes by 0A∼P for each strain is calculated in the same way as described in Section I, *Eqn. I. 6, Eqn. I. 8*, and *Eqn. I. 9* for DNA fragments with one, two, and three boxes, respectively, using the best-fit binding energy and interaction energy parameters shown in Supplementary Table S4. Occupancy probability for each mutant strain is shown in Supplementary Figure S6.

Notably, the expressions representing the Boltzmann weights of different 0A∼P bound configurations in *Eqn. I. 6, Eqn. I. 8*, and *Eqn. I. 9* also appear in the calculation of the Boltzmann weight in *Eqn. II. 4*, in both promoter-bound and promoter-unbound configurations. If there is no time-dependent change in RNAP concentration, and given that the maximum transcription initiation rate is constant over all time points, there is no change in effective transcription rate in accordance with *Eqn. II. 2*, i.e., saturated 0A∼P binding probability will result in saturated effective transcription rate.

### (VII) Bayesian Information Criterion for differentially constrained models

To verify that the model is not overfitted upon the addition of interaction energy parameters, we computed the Bayesian Information Criterion (11) (BIC) for the models with none or up to three secondary interaction energies (*G*_12_, *G*_13_, *G*_23_) being zero.

To compute the BIC, we use the following formula (11):

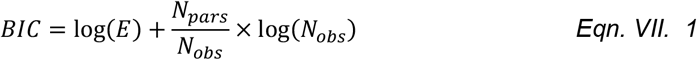

Where *E* is the error of the objective function employed during optimization of the biophysical model (Methods and Section 4.5). The variable *N*_*pars*_ is the number of unknown fitting parameters, and *N*_*obs*_ is the total number of experimental data points.

The lowest BIC is found in the model with all secondary interaction energies being nonzero (Supplementary Figure Table S5). As such, the model with the existence of higher-order interaction energies is not overfitting the experimental data, and the kinetic control model requires the attractions between bound 0A boxes to explain *spo0A* autoregulation.

### (VIII) Model-predicted regulatory role of 0A boxes in *v*_*max*_ and *v*^*effective*^

In Figure 6, we examined the model-predicted effect of 0A∼P binding at different boxes in two ways: 1) by analyzing the change in maximum transcription initiation rate of each configuration (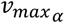, Figure 6A-C) relative to the configuration ‘000’; 2) by analyzing the effective transcription rate at t = 9 h of each mutant strain (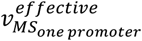 and 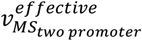, Figure 6D) relative to the ‘none’ strain. Below, we show that these two tests are analytically equivalent for a single-promoter kinetic control model. We then show that, for the double-promoter kinetic control model, the two tests are not equivalent, as the 0A∼P binding effect on the dual-promoter effective transcription rate is the weighted sum of 0A∼P binding effects on *Pv* and *Ps*, respectively.

#### (viii.1) for single-promoter strains

Here, we will demonstrate that the relative 0A box effects on *v*_*max*_ and *v*^*effective*^ are quantitatively equivalent in single-promoter strains. We assume the most complex case with all 0A boxes being present (‘123 strain’). Due to the nature of the kinetic control model, in accordance with *Eqn. IV. 12*, the effective transcription rate can be written as a product of the factorization term (denoting RNAP binding) and the sum of 0A∼P binding weighted by configuration-dependent maximum transcription initiation rate:

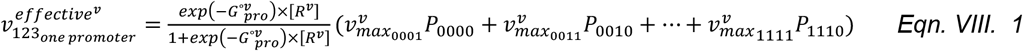

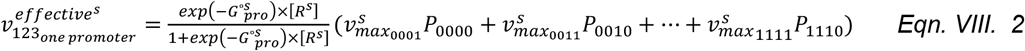

Here, superscript ‘v’ or ‘s’ are promoter-specific parameters. The binary subscripts for *v*_*max*_ are written in terms of promoter-configurations *α*_*T*_; the binary subscript for *P* are written in terms of promoter-unbound configurations *α*_*TF*_, in accordance with Section iv.2.a and *Eqn. IV. 12*. For example, ‘0011’ denotes 0A3 box bound and promoter bound.

At t = 9 h, for strain ‘123’, the probability of all boxes being bound (P_1110_) is equal to 1 (Figure S6). As such, all other possible configurations have a probability equal to zero. Given this, the effective transcription rate for P*v* and P*s* ‘123’ can be simplified into:

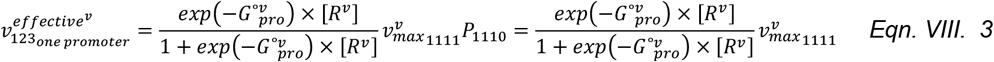

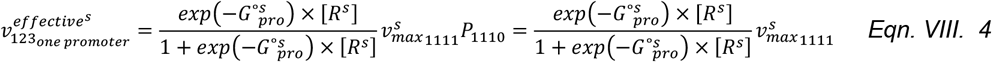

In Figure 6D, the effective transcription rate of each strain is relative to the ‘none’ strain. Using *Eqn. II. 2, Eqn. II. 5* and *Eqn. II. 6*, the effective transcription rate for ‘none’ strain can be written as below, with [0*A*∼*P*] = 0 and *α*_*T*_ = [0,0,0,1]:

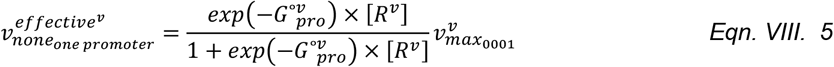

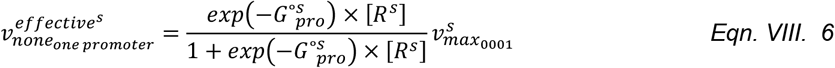

The fold change in ‘123’ strain compared to ‘none’ strain, therefore, is the ratio between *Eqn. VIII. 3 and Eqn. VIII. 5*, and the ratio between *Eqn. VIII. 4 and Eqn. VIII. 6*, for P*v* and P*s*, respectively:

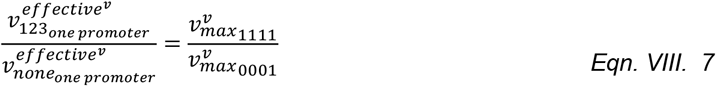

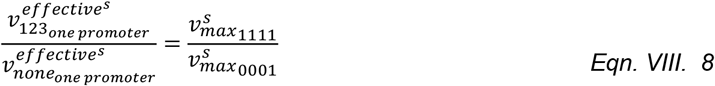

Here, the fold change in effective transcription rate between ‘123’ and ‘none’ strain is the same as normalizing *v*_*max*_ of configuration ‘111’ (where maximum number of allowable boxes are bound in ‘123’) by the *v*_*max*_ of configuration ‘000’. This is true for all other mutant strains, because of the saturated effect of 0A∼P binding at t = 9h for all mutant strains. Moreover, the fold change does not depend on the concentration of RNAP.

#### (viii.2) for double-promoter strains

Using the additive property of the kinetic control model (*Eqn. IV. 25*) and knowing that 0A∼P binding probability is contributed exclusively by a single configuration, which saturates at the last time point (as shown in section viii.1), the effective transcription rate for ‘123’ and ‘none’ for ‘P*v*P*s* strains’ can be directly represented using *Eqn. VIII. 3-6*:

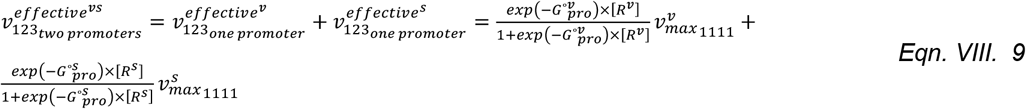

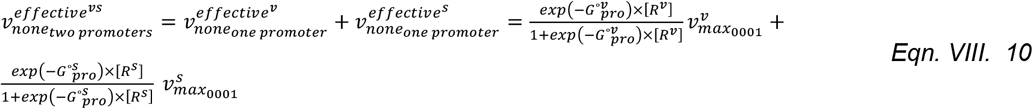

For convenience, we can define

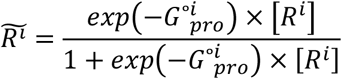

with i ∈ {*s, v*}.

When computing the ratio between ‘123’ and ‘none’ for ‘*PvPs* strains’, we obtain the following expression:

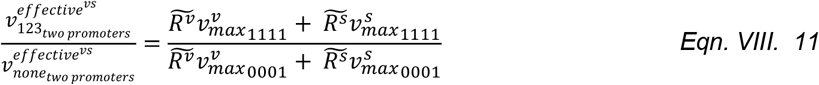

To further simplify this equation, we divide the top and bottom of *Eqn. VIII. 11* by 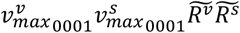. In doing so, we obtain:

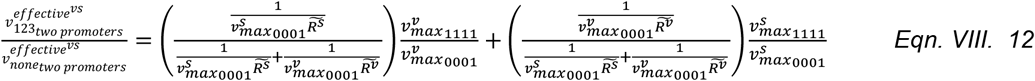

Here, we observe that the effect of 0A∼P binding on a two-promoter strain is the weighted sum of the effect on the single-promoter expression, i.e., the weighted sum of the ratio between P*v*-bound and P*s*-bound *v*_*max*_ (weighted sum of *Eqn. VIII. 7-8*). In addition, the fold change of the ‘123’ strain in relative to ‘none’ strain now depends on the concentration of RNAP.

### (XI) Find RNAP holoenzyme dynamics common to *spo0A* and non-*spo0A* promoters

In Result Section 2.5, to verify whether *σ*^A^- and *σ*^H^-RNAP holoenzyme levels are time-dependent in general, we measured the activity of two non-*spo0A* promoters: P*hyper-spac* (*σ*^A^ specific) and *citG* promoter P*citG* (*σ*^H^ specific) over 7 hours. Below we show the method adopted to find a common time-dependent *σ*^A^ -RNAP increase that explains both P*hyper-spac* and P*v* gene expression. The same method can be applied to finding the *σ*^H^-RNAP increase for P*citG* and P*s*.

To start with, we assume the increase in *σ*^A^-RNAP is identical between the ‘P*v* strain, none’ expression and the constitutive expression of P*hyper-spac*, both of which do not depend on additional transcription factor binding (i.e., ‘none’ strain). Based on the same modeling framework for a single promoter (Supplementary Section II), there are only two binding configurations possible for such DNA fragment with one promoter (i.e., [*σ*_1_ = 0, *σ*_2_ = 0, *σ*_3_ = 0, *σ*_*pro*_ = 0] and [*σ*_1_ = 0, *σ*_2_ = 0, *σ*_3_ = 0, *σ*_*pro*_ = 1]). Thus, the effective transcription rate of each promoter, contributed by the promoter-bound configuration, can be written using *Eqn. II. 2*:

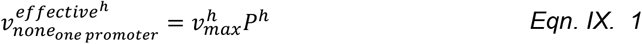

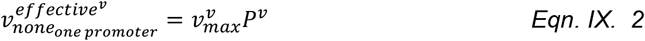

Here, the superscript ‘h’ or ‘v’ denotes promoter-specific maximum transcription initiation rate and promoter-specific RNAP binding probability, for the configuration [*σ*_1_ = 0, *σ*_2_ = 0, *σ*_3_ = 0, *σ*_*pro*_ = 1] in each promoter.

The RNAP binding probability of each promoter can be written using *Eqn. II. 5* and *Eqn. II. 6* as:

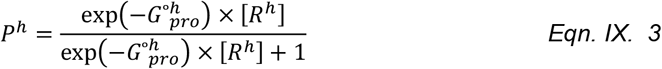

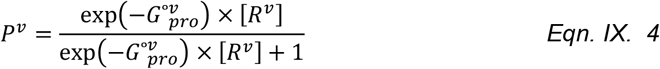

where, 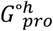 and 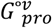 are RNAP binding affinity to the *hyspac and spo0A* vegetative promoter, respectively. Here, there is no binding of transcription factor such as 0A∼P (i.e., [0*A*∼*P*] = 0).

In accordance with the assumption in the RNAP holoenzyme, we rename the RNAP concentration variable as [*R*^*common*^]:

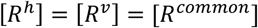

By examining *Eqn. IX. 1-4*, one can see that there are two parameters (i.e., *v*_*max*_ and *G*_*pro*_) that can be potentially responsible for the difference in effective transcription rate between P*hyspac* and P*v (*i.e., between 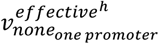 and 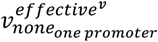). Here, we assume that the maximum transcription initiation rate is the same between the promoters (renamed as 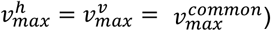, but the binding affinity of each promoter is unique 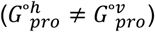.

To this end, let the gene expression of P*hyspac* and P*v* at a particular time point *t* = *w* hour be 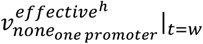 and 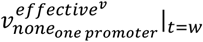, and let the RNAP holoenzyme concentration at the same time point be [*R*]_*t*=*w*_. As such, the ratio of model-predicted effective transcription rate between two promoters at *t* = *w* hour can be expressed as:

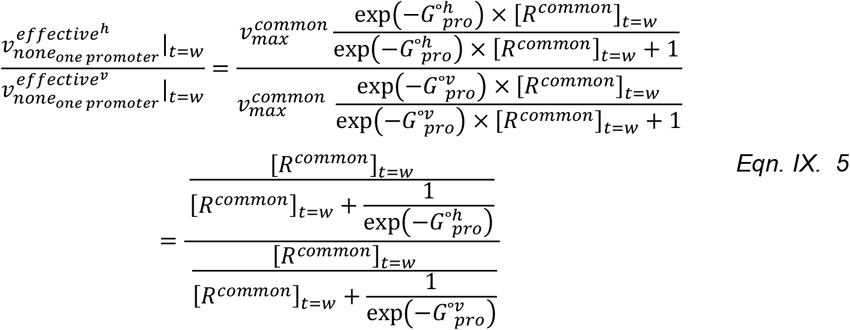

Next, we define the ratio in experimentally measured promoter activity between two promoters at *t* = *w* hour as 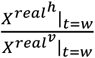

To find the *σ*^A^-RNAP increase that explains both P*hyper-spac* and P*v* gene expression, we minimize the difference (*E*_*sigma*_) between model-predicted and measured ratio of gene expression across all time points:

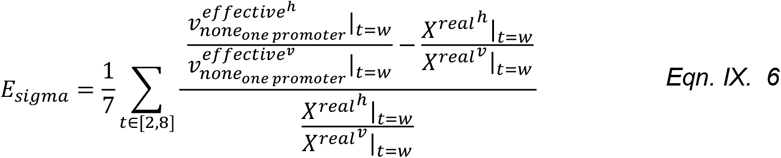

We found reasonable 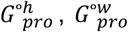, and [*R*^*common*^]_*t*=*w*_ (*i* ∈ [2,8]) parameters using MATLAB (2021a) Particle Swarm Optimization (2).

## Supplementary Tables

**Table S1.** List of *lacZ* reporter fusion strains used. Also shown is the ‘Alias’ name used to refer to each strain in the main manuscript and supplementary files.

| *B. subtilis* strains identifier | Reference | Genotype | Alias |
| --- | --- | --- | --- |
| amyE::Pspo0A<sup>wt</sup>-lacZ spc coml<sup>Q12L</sup> (9766 → 5609) S | (1) | No Pv or Ps removal, no 0A mutation | PvPs strain, 123 |
| amyE::Pspo0A<sup>0A1*</sup>-lacZ spc coml<sup>Q12L</sup> (9673 → 5609) S | (1) | No Pv or Ps removal, 0A1 mutated | PvPs strain, 23 |
| amyE::Pspo0A<sup>0A2*</sup>-lacZ spc coml<sup>Q12L</sup> (9674 → 5609) S | (1) | No Pv or Ps removal, 0A2 mutated | PvPs strain, 13 |
| amyE::Pspo0A<sup>0A3*</sup>-lacZ spc coml<sup>Q12L</sup> (9675 → 5609) S | (1) | No Pv or Ps removal, 0A3 mutated | PvPs strain, 12 |
| amyE::Pspo0A<sup>0A1*2*</sup>-lacZ spc coml<sup>Q12L</sup> (pMF1056 → 5609) S | (1) | No Pv or Ps removal, 0A1 and 0A2 mutated | PvPs strain, 3 |
| amyE::Pspo0A<sup>0A1*3*</sup>-lacZ spc coml<sup>Q12L</sup> (pMF1010 → 5609) S | (1) | No Pv or Ps removal, 0A1 and 0A3 mutated | PvPs strain, 2 |
| amyE::Pspo0A<sup>0A2*3*</sup>-lacZ spc coml<sup>Q12L</sup> (pMF1057 → 5609) S | (1) | No Pv or Ps removal, 0A2 and 0A3 mutated | PvPs strain, 1 |
| amyE::Pspo0A<sup>0A1*2*3*</sup>-lacZ spc coml<sup>Q12L</sup> (pMF1058 → 5609) S | (1) | No Pv or Ps removal, 0A1, 0A2, 0A3 mutated | PvPs strain, none |
| amyE::Pspo0A<sup>0A4*</sup>-lacZ spc coml<sup>Q12L</sup> (10128 → 5609) S | (1) | Ps removed, no 0A mutation | Pv strain, 123 |
| amyE::Pspo0A<sup>0A1*4*</sup>-lacZ spc coml<sup>Q12L</sup> (10129 → 5609) S | (1) | Ps removed, 0A1 mutated | Pv strain, 23 |
| amyE::Pspo0A<sup>0A2*4*</sup>-lacZ spc coml<sup>Q12L</sup> (10130 → 5609) S | (1) | Ps removed, 0A2 mutated | Pv strain, 13 |
| amyE::Pspo0A<sup>0A3*4*</sup>-lacZ spc coml<sup>Q12L</sup> (10131 → 5609) S | (1) | Ps removed, 0A3 mutated | Pv strain, 12 |
| amyE::Pspo0A<sup>0A1*2*4*</sup>-lacZ spc coml<sup>Q12L</sup> (pMF1059 → 5609) S | (1) | Ps removed, 0A1 and 0A2 mutated | Pv strain, 3 |
| amyE::Pspo0A<sup>0A1*3*4*</sup>-lacZ spc coml<sup>Q12L</sup> (pMF1043 → 5609) S | (1) | Ps removed, 0A1 and 0A3 mutated | Pv strain, 2 |
| amyE::Pspo0A<sup>0A2*3*4*</sup>-lacZ spc coml<sup>Q12L</sup> (pMF1060 → 5609) S | (1) | Ps removed, 0A2 and 0A3 mutated | Pv strain, 1 |
| amyE::P<sub>spo0A</sub><sup>0A1*2*3*4*</sup>-lacZ spc comI<sup>Q12L</sup> (pMF1061 → 5609) S | (1) | Ps removed, 0A1, 0A2, 0A3 mutated | Pv strain, none |
| amyE::P<sub>Δv</sub><sup>wt</sup>-lacZ spc comI<sup>Q12L</sup> (9795 → 5609) S | (1) | Pv removed, no 0A mutation | Ps strain, 123 |
| amyE::P<sub>Δv</sub><sup>0A1*</sup>-lacZ spc comI<sup>Q12L</sup> (pMF1068 → 5609) S | (1) | Pv removed, 0A1 mutated | Ps strain, 23 |
| amyE::P<sub>Δv</sub><sup>0A2*</sup>-lacZ spc comI<sup>Q12L</sup> (9796 → 5609) S | (1) | Pv removed, 0A2 mutated | Ps strain, 13 |
| amyE::P<sub>Δv</sub><sup>0A3*</sup>-lacZ spc comI<sup>Q12L</sup> (9797 → 5609) S | (1) | Pv removed, 0A3 mutated | Ps strain, 12 |
| amyE::P<sub>Δv</sub><sup>0A1*2*</sup>-lacZ spc comI<sup>Q12L</sup> (pMF1070 → 5609) S | (1) | Pv removed, 0A1 and 0A2 mutated | Ps strain, 3 |
| amyE::P<sub>Δv</sub><sup>0A1*3*</sup>-lacZ spc comI<sup>Q12L</sup> (pMF1071 → 5609) S | (1) | Pv removed, 0A1 and 0A3 mutated | Ps strain, 2 |
| amyE::P<sub>Δv</sub><sup>0A2*3*</sup>-lacZ spc comI<sup>Q12L</sup> (pMF1072 → 5609) S | (1) | Pv removed, 0A2 and 0A3 mutated | Ps strain, 1 |
| amyE::P<sub>Δv</sub><sup>0A1*2*3*</sup>-lacZ spc comI<sup>Q12L</sup> (pMF1073 → 5609) S | (1) | Pv removed, 0A1, 0A2, 0A3 mutated | Ps strain, none |
| amyE::Phy-spac-lacZ cm comI<sup>Q12L</sup> (342 → 5609) S | Original to this study | Constitutive σ<sup>A</sup> specific *hyper-spac* promoter | *Phyper-spac* |
| amyE::PcitG-lacZ spc comI<sup>Q12L</sup> (877 → 5609) C | Original to this study | Constitutive σ<sup>H</sup> specific *citG* promoter | *PcitG* |

**Table S2.** Optimization error of the thermodynamic model fit to EMSA data under different assumptions of cooperativity. The fold change in optimization error (Supplementary Section I, subsection i.3) between the two optimizations with different *N* is ∼3.7 fold.

| Cooperativity | Optimization error |
| --- | --- |
| 2 | 0.1410 |
| 4 | 0.0382 |

**Table S3.** List of the parameters used in the biophysical models.

| Parameters | Description |
| --- | --- |
| $G^\circ_1$ | Binding energy of 0A~P to 0A1 |
| $G^\circ_2$ | Binding energy of 0A~P to 0A2 |
| $G^\circ_3$ | Binding energy of 0A~P to 0A3 |
| $G^\circ_v$ | Binding energy of RNAP to $P_v$ promoter |
| $G^\circ_s$ | Binding energy of RNAP to $P_s$ promoter |
| $G^\circ_{12}$ | Interaction energy between 0A~P bound at 0A1 and 0A2 |
| $G^\circ_{13}$ | Interaction energy between 0A~P bound at 0A1 and 0A3 |
| $G^\circ_{23}$ | Interaction energy between 0A~P bound at 0A2 and 0A3 |
| $G^\circ_{123}$ | Interaction energy between 0A~P bound at 0A1, 0A2, and 0A3 |
| $[0A\sim P]_{t=w}$ | Concentration of 0A~P at the $w^{\text{th}}$ hour. $w \in [2,9]$ |
| $[R^s]_{t=w}$ | Concentration of sigma H-associated RNAP at the $w^{\text{th}}$ hour. $w \in [2,9]$ |
| $[R^v]_{t=w}$ | Concentration of sigma A-associated RNAP at the $w^{\text{th}}$ hour. $w \in [2,9]$ |
| $G_{1pro}^{\circ v}$ | Interaction energy between 0A~P bound at 0A1 and RNAP bound at $Pv$ |
| $G_{2pro}^{\circ v}$ | Interaction energy between 0A~P bound at 0A2 and RNAP bound at $Pv$ |
| $G_{3pro}^{\circ v}$ | Interaction energy between 0A~P bound at 0A3 and RNAP bound at $Pv$ |
| $G_{12pro}^{\circ v}$ | Interaction energy between 0A~P bound at 0A1, 0A~P bound at 0A2, and RNAP bound at $Pv$ |
| $G_{13pro}^{\circ v}$ | Interaction energy between 0A~P bound at 0A1, 0A~P bound at 0A3, and RNAP bound at $Pv$ |
| $G_{23pro}^{\circ v}$ | Interaction energy between 0A~P bound at 0A2, 0A~P bound at 0A3, and RNAP bound at $Pv$ |
| $G_{123pro}^{\circ v}$ | Interaction energy between 0A~P bound at 0A1, 0A~P bound at 0A2, 0A~P bound at 0A3, and RNAP bound at $Pv$ |
| $v_{max_{0001}}^v$ | Maximum transcription initiation rate for configurations where $Pv$ is bound but no 0A box is bound |
| $v_{max_{0011}}^v$ | Maximum transcription initiation rate for configurations where $Pv$ is bound and 0A3 is bound |
| $v_{max_{0101}}^v$ | Maximum transcription initiation rate for configurations where $Pv$ is bound and 0A2 is bound |
| $v_{max_{0111}}^v$ | Maximum transcription initiation rate for configurations where $Pv$ is bound, 0A2 and 0A3 are bound |
| $v_{max_{1001}}^v$ | Maximum transcription initiation rate for configurations where $Pv$ is bound and 0A1 is bound |
| $v_{max_{1011}}^v$ | Maximum transcription initiation rate for configurations where $Pv$ is bound, and 0A1 and 0A3 are bound |
| $v_{max_{1101}}^v$ | Maximum transcription initiation rate for configurations where $Pv$ is bound, and 0A1 and 0A2 are bound |
| $v_{max_{1111}}^v$ | Maximum transcription initiation rate for configurations where $Pv$ is bound, and 0A1, 0A2, and 0A3 are bound |
| $G_{1pro}^{os}$ | Interaction energy between 0A~P bound at 0A1 and RNAP bound at $Ps$ |
| $G_{2pro}^{os}$ | Interaction energy between 0A~P bound at 0A2 and RNAP bound at $Ps$ |
| $G_{3pro}^{os}$ | Interaction energy between 0A~P bound at 0A3 and RNAP bound at $Ps$ |
| $G_{12pro}^{os}$ | Interaction energy between 0A~P bound at 0A1, 0A~P bound at 0A2, and RNAP bound at $Ps$ |
| $G_{13pro}^{os}$ | Interaction energy between 0A~P bound at 0A1, 0A~P bound at 0A3, and RNAP bound at $Ps$ |
| $G_{23pro}^{os}$ | Interaction energy between 0A~P bound at 0A2, 0A~P bound at 0A3, and RNAP bound at $Ps$ |
| $G_{123pro}^{os}$ | Interaction energy between 0A~P bound at 0A1, 0A~P bound at 0A2, 0A~P bound at 0A3, and RNAP bound at $Ps$ |
| $v_{max_{0001}}^s$ | Maximum transcription initiation rate for configurations where $Ps$ is bound but no 0A box is bound |
| $v_{max_{0011}}^s$ | Maximum transcription initiation rate for configurations where $Ps$ is bound and 0A3 is bound |
| $v_{max_{0101}}^s$ | Maximum transcription initiation rate for configurations where $Ps$ is bound and 0A2 is bound |
| $v_{max_{0111}}^s$ | Maximum transcription initiation rate for configurations where $Ps$ is bound and 0A2 and 0A3 are bound |
| $v_{max_{1001}}^s$ | Maximum transcription initiation rate for configurations where $Ps$ is bound and 0A1 are bound |
| $v_{max_{1011}}^s$ | Maximum transcription initiation rate for configurations where $Ps$ is bound, and 0A1 and 0A3 are bound |
| $v_{max_{1101}}^s$ | Maximum transcription initiation rate for configurations where $Ps$ is bound, and 0A1 and 0A2 are bound |
| $v_{max_{1111}}^s$ | Maximum transcription initiation rate for configurations where $Ps$ is bound, and 0A1, 0A2, and 0A3 are bound |

**Table S4.** Model-predicted parameter values from the best fit purely kinetic model with existence of all higher-order interaction energies. Higher-order interaction energies refer to *G*°_12_, *G*°_13_, *G*°_23_, and *G*°_123_ in the biophysical model. The mean of predicted parameters across 20 independent optimization runs is reported with standard deviation. Full parameter list and corresponding descriptions are shown in Table S3. Note that energies between 0A boxes and promoters do not exist (values are zero, thus not shown) in the purely kinetic biophysical model.

| Parameter | Mean | Std | Unit |
| --- | --- | --- | --- |
| $G_1^{\circ}$ | -5.42 | <10e-3 | kT |
| $G_2^{\circ}$ | -2.35 | <10e-3 | kT |
| $G_3^{\circ}$ | -7.40 | <10e-3 | kT |
| $G_{12}^{\circ}$ | -8.54 | 1.22 | kT |
| $G_{13}^{\circ}$ | -8.82 | 5.04 | kT |
| $G_{23}^{\circ}$ | -13.96 | 0.087 | kT |
| $G_{123}^{\circ}$ | -9.58 | 7.0283 | kT |
| $v_{max_{0001}}^v$ | 685.03 | 22.17 | a.u. |
| $v_{max_{0011}}^v$ | 617.18 | 13.80 | a.u. |
| $v_{max_{0101}}^v$ | 628.38 | 11.17 | a.u. |
| $v_{max_{0111}}^v$ | 693.85 | 17.22 | a.u. |
| $v_{max_{1001}}^v$ | 410.64 | 7.77 | a.u. |
| $v_{max_{1011}}^v$ | 412.87 | 11.94 | a.u. |
| $v_{max_{1101}}^v$ | 350.20 | 5.93 | a.u. |
| $v_{max_{1111}}^v$ | 466.28 | 14.93 | a.u. |
| $v_{max_{0001}}^S$ | 977.09 | 34.19 | a.u. |
| $v_{max_{0011}}^S$ | 1254.91 | 41.21 | a.u. |
| $v_{max_{0101}}^S$ | 736.31 | 23.62 | a.u. |
| $v_{max_{0111}}^S$ | 149.61 | 5.15 | a.u. |
| $v_{max_{1001}}^S$ | 1077.36 | 34.68 | a.u. |
| $v_{max_{1011}}^S$ | 1233.93 | 40.93 | a.u. |
| $v_{max_{1101}}^S$ | 680.73 | 23.31 | a.u. |
| $v_{max_{1111}}^S$ | 280.15 | 10.18 | a.u. |
| $[R^v]_{t=2}$ | 0.25 | 0.01 | $\mu\text{M}$ |
| $[R^v]_{t=3}$ | 0.35 | 0.02 | $\mu\text{M}$ |
| $[R^v]_{t=4}$ | 0.35 | 0.01 | $\mu\text{M}$ |
| $[R^v]_{t=5}$ | 0.50 | 0.02 | $\mu\text{M}$ |
| $[R^v]_{t=6}$ | 0.73 | 0.03 | $\mu\text{M}$ |
| $[R^v]_{t=7}$ | 1.11 | 0.06 | $\mu\text{M}$ |
| $[R^v]_{t=8}$ | 1.37 | 0.08 | $\mu\text{M}$ |
| $[R^v]_{t=9}$ | 2.00 | 0.02 | $\mu\text{M}$ |
| $[R^s]_{t=2}$ | 0.06 | 0.002 | $\mu\text{M}$ |
| $[R^s]_{t=3}$ | 0.13 | 0.005 | $\mu\text{M}$ |
| $[R^s]_{t=4}$ | 0.44 | 0.02 | $\mu\text{M}$ |
| $[R^s]_{t=5}$ | 0.73 | 0.05 | $\mu\text{M}$ |
| $[R^s]_{t=6}$ | 1.08 | 0.07 | $\mu\text{M}$ |
| $[R^S]_{t=7}$ | 1.33 | 0.09 | $\mu\text{M}$ |
| $[R^S]_{t=8}$ | 1.73 | 0.14 | $\mu\text{M}$ |
| $[R^S]_{t=9}$ | 1.90 | 0.15 | $\mu\text{M}$ |

**Table S5.** The optimization error of the purely kinetic biophysical model under different assumptions on the existence of high-order interaction energy parameters. The optimization error is computed based on Material and Methods Section 4.5, *Eqn. 5*. Higher-order interaction energies refer to *G*°_12_, *G*°_13_, *G*°_23_, and *G*°_123_ in the biophysical model. Shown in the second column is, under the current assumption, how many higher-order interaction energies are nonzero (i.e., the energy exists). The Bayesian information criterion (BIC) was computed for each assumption. In grey is the best fitting model (i.e. lowest optimization error), where all secondary energy parameters are nonzero and tertiary energy parameter is zero. Full parameter list and descriptions are shown in Table S3.

| Assumption for energy parameters | Number of higher-order energies being nonzero | Optimization error | BIC |
| --- | --- | --- | --- |
| $G_{12}^\circ = 0, G_{13}^\circ = 0, G_{23}^\circ = 0, G_{123}^\circ = 0$ | 0 | 0.1886 | -0.7646 |
| $G_{12}^\circ \neq 0, G_{13}^\circ = 0, G_{23}^\circ = 0, G_{123}^\circ = 0$ | 1 | 0.1665 | -0.8619 |
| $G_{12}^\circ = 0, G_{13}^\circ \neq 0, G_{23}^\circ = 0, G_{123}^\circ = 0$ | 1 | 0.1807 | -0.7799 |
| $G_{12}^\circ = 0, G_{13}^\circ = 0, G_{23}^\circ \neq 0, G_{123}^\circ = 0$ | 1 | 0.1375 | -1.0533 |
| $G_{12}^\circ = 0, G_{13}^\circ = 0, G_{23}^\circ = 0, G_{123}^\circ \neq 0$ | 1 | 0.1740 | -0.8176 |
| $G_{12}^\circ \neq 0, G_{13}^\circ \neq 0, G_{23}^\circ = 0, G_{123}^\circ = 0$ | 2 | 0.1538 | -0.9140 |
| $G_{12}^{\circ} \neq 0, G_{13}^{\circ} = 0, G_{23}^{\circ} \neq 0, G_{123}^{\circ} = 0$ | 2 | 0.1118 | -1.1720 |
| $G_{12}^{\circ} \neq 0, G_{13}^{\circ} = 0, G_{23}^{\circ} = 0, G_{123}^{\circ} \neq 0$ | 2 | 0.1508 | -0.9333 |
| $G_{12}^{\circ} = 0, G_{13}^{\circ} \neq 0, G_{23}^{\circ} \neq 0, G_{123}^{\circ} = 0$ | 2 | 0.1286 | -1.0929 |
| $G_{12}^{\circ} = 0, G_{13}^{\circ} \neq 0, G_{23}^{\circ} = 0, G_{123}^{\circ} \neq 0$ | 2 | 0.1605 | -0.8708 |
| $G_{12}^{\circ} = 0, G_{13}^{\circ} = 0, G_{23}^{\circ} \neq 0, G_{123}^{\circ} \neq 0$ | 2 | 0.1259 | -1.1140 |
| $G_{12}^{\circ} \neq 0, G_{13}^{\circ} \neq 0, G_{23}^{\circ} \neq 0, G_{123}^{\circ} = 0$ | 3 | 0.0731 | -1.6301 |
| $G_{12}^{\circ} \neq 0, G_{13}^{\circ} \neq 0, G_{23}^{\circ} = 0, G_{123}^{\circ} \neq 0$ | 3 | 0.1369 | -1.0026 |
| $G_{12}^{\circ} \neq 0, G_{13}^{\circ} = 0, G_{23}^{\circ} \neq 0, G_{123}^{\circ} \neq 0$ | 3 | 0.0774 | -1.5725 |
| $G_{12}^{\circ} = 0, G_{13}^{\circ} \neq 0, G_{23}^{\circ} \neq 0, G_{123}^{\circ} \neq 0$ | 3 | 0.0858 | -1.4697 |
| $G_{12}^{\circ} \neq 0, G_{13}^{\circ} \neq 0, G_{23}^{\circ} \neq 0, G_{123}^{\circ} \neq 0$ | 4 | 0.0731 | -1.6033 |

**Table S6.**
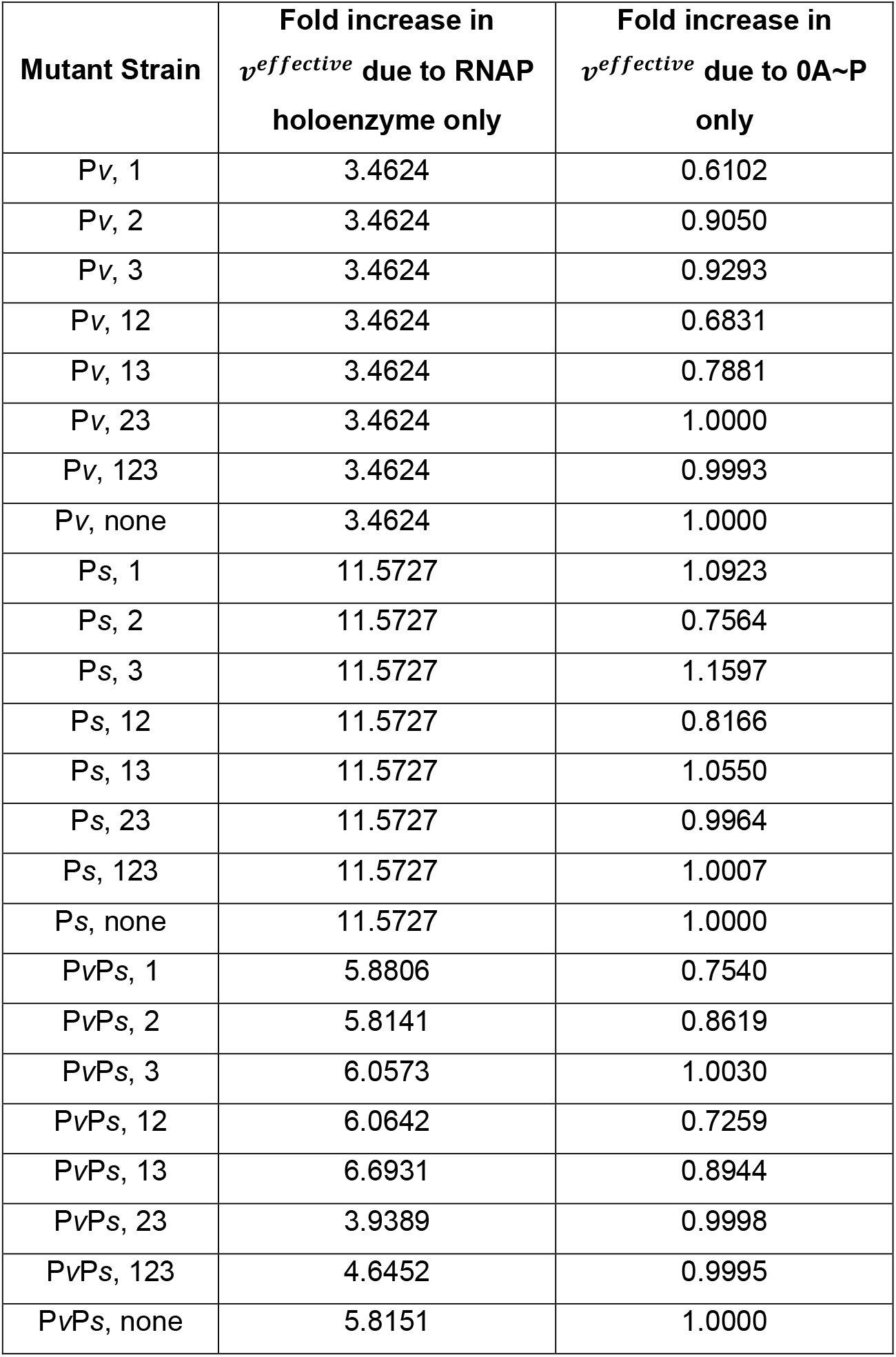
Dissection of the effect of 0A∼P binding and RNAP binding on *spo0A* expression for each mutant strain. Using best-fit model parameters (Table S4), the biophysical model was simulated with either constant 0A∼P concentration (second column) or constant RNAP holoenzyme concentration (third column) over time. For each simulation, the fold change of effective transcription rate in the last time point (t = 9h) in comparison to the first time point (t = 2h) was computed for each entry.

| **Mutant Strain** | **Fold increase in $v^{effective}$ due to RNAP holoenzyme only** | **Fold increase in $v^{effective}$ due to 0A~P only** |
| --- | --- | --- |
| Pv, 1 | 3.4624 | 0.6102 |
| Pv, 2 | 3.4624 | 0.9050 |
| Pv, 3 | 3.4624 | 0.9293 |
| Pv, 12 | 3.4624 | 0.6831 |
| Pv, 13 | 3.4624 | 0.7881 |
| Pv, 23 | 3.4624 | 1.0000 |
| Pv, 123 | 3.4624 | 0.9993 |
| Pv, none | 3.4624 | 1.0000 |
| Ps, 1 | 11.5727 | 1.0923 |
| Ps, 2 | 11.5727 | 0.7564 |
| Ps, 3 | 11.5727 | 1.1597 |
| Ps, 12 | 11.5727 | 0.8166 |
| Ps, 13 | 11.5727 | 1.0550 |
| Ps, 23 | 11.5727 | 0.9964 |
| Ps, 123 | 11.5727 | 1.0007 |
| Ps, none | 11.5727 | 1.0000 |
| PvPs, 1 | 5.8806 | 0.7540 |
| PvPs, 2 | 5.8141 | 0.8619 |
| PvPs, 3 | 6.0573 | 1.0030 |
| PvPs, 12 | 6.0642 | 0.7259 |
| PvPs, 13 | 6.6931 | 0.8944 |
| PvPs, 23 | 3.9389 | 0.9998 |
| PvPs, 123 | 4.6452 | 0.9995 |
| PvPs, none | 5.8151 | 1.0000 |

## Supplementary Figures

**Figure S1.**
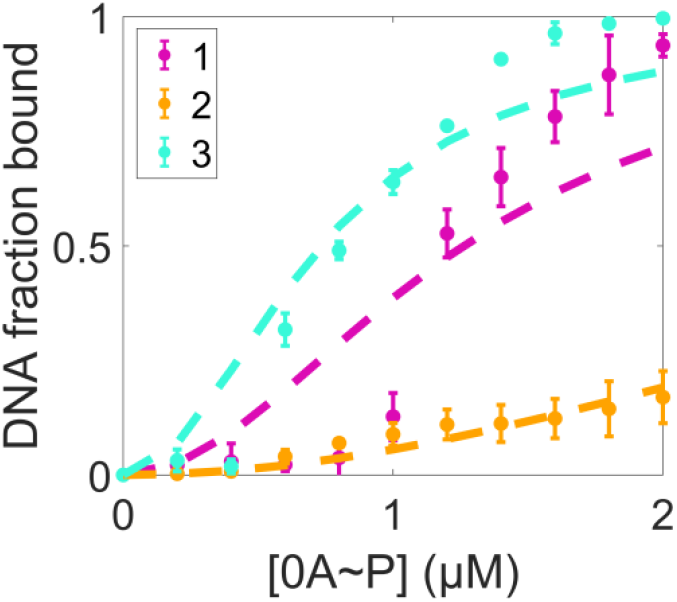
Fitting the experimentally determined fraction of bound DNA to a statistical thermodynamic model with Hill coefficient (*N*) of 2. Model prediction (dashed lines) and empirical measurement (solid lines with standard deviation error bars) are shown for DNA fragments with one 0A box mutation. The fitting is worse than when considering cooperativity (*N*) of 4.

**Figure S2.**
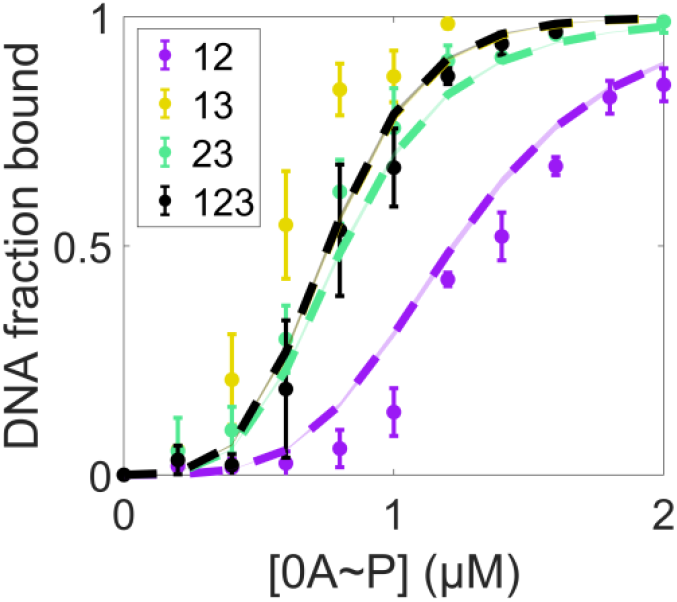
Predicting fraction of bound DNA for constructs with multiple boxes present using single binding energies optimized from statistical thermodynamic model. Solid data points with error bars are experimental measurement with standard deviation. Dashed lines with colored patches are mean predictions with standard deviations, using different parameter sets derived from 20 augmented measurements. Dashed yellow and dashed black lines overlap.

**Figure S3.**
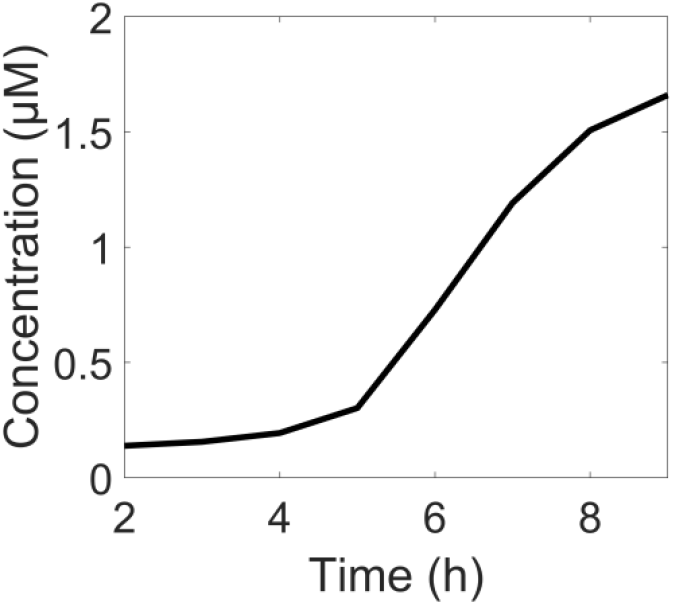
Model input dynamics of Spo0A∼P concentration for the wildtype (WT) strain. Predicted dynamics are based on (12).

**Figure S4.**
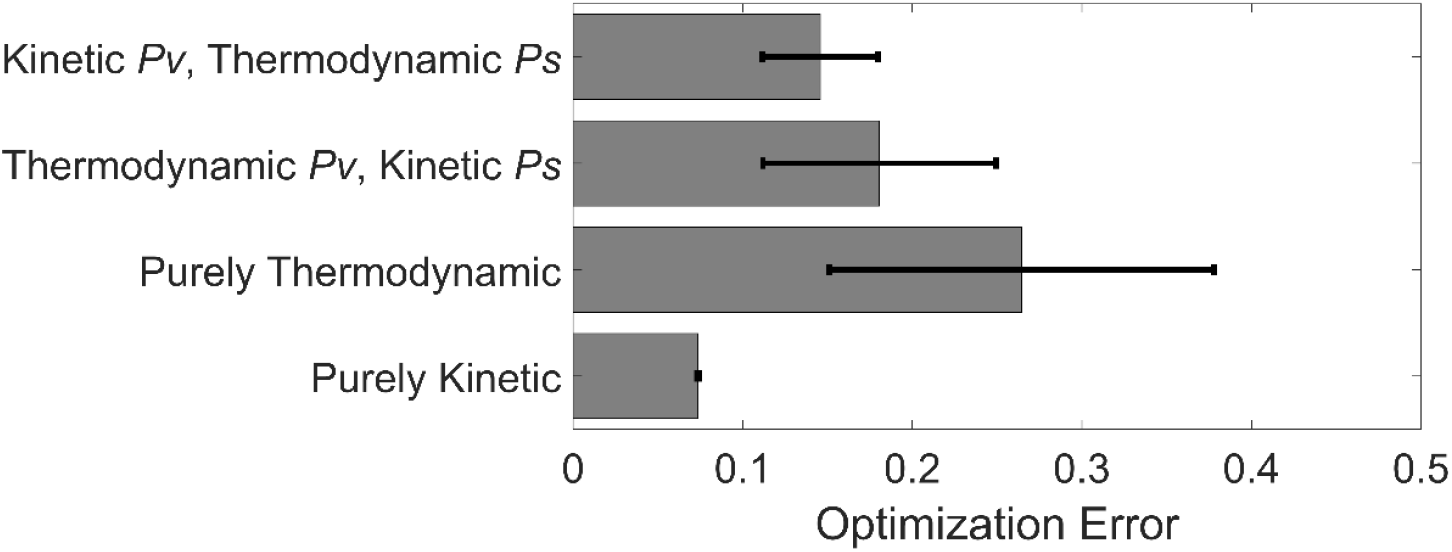
Optimization error for all biophysical models examined. Each model was optimized at least 20 times to examine if it best-fits the experimental *spo0A* promoter activity using MATLAB Particle Swarm Optimization algorithm (2). The number of fitting parameters were kept the same across all models. Shown is the mean and standard deviation of the output of the objective function (Material and Methods Section 4.5, *Eqn. 5*) for the four transcriptional control models tested.

**Figure S5.**
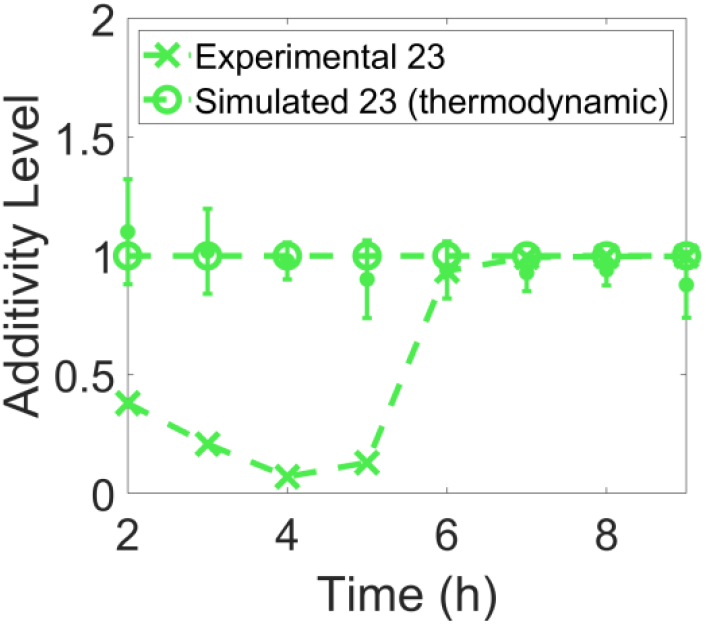
Additivity of thermodynamic and kinetic control models. Additivity level for the purely kinetic model (marker “O”) and for the purely thermodynamic model (marker “X”) with two 0A boxes on the DNA fragment, under randomly sampled energy parameters. The empirical data (solid line with colored regions and error bar) with two 0A boxes (0A2 and 0A3) are shown for reference.

**Figure S6.**
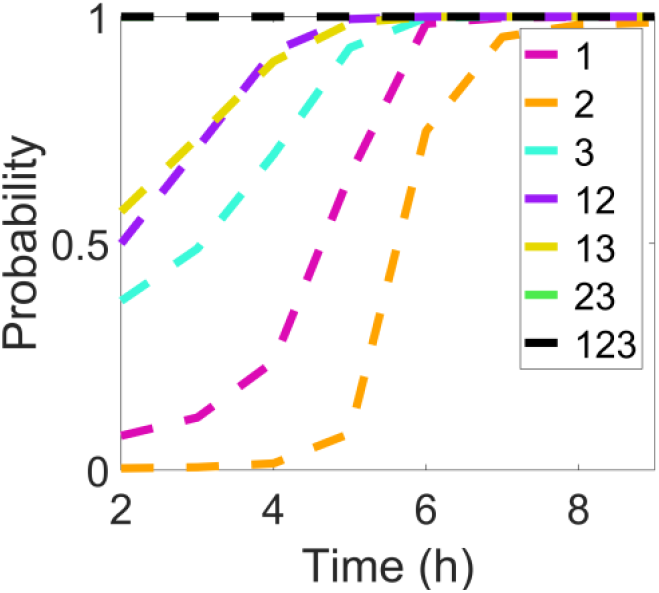
Probability that all 0A boxes are bound for strains with one, two, or three boxes on the DNA fragment. Occupancy probability is computed with 0A∼P dynamics shown in Supplementary Figure S3 and best-fit energy parameters in Supplementary Table S4.

**Figure S7.**
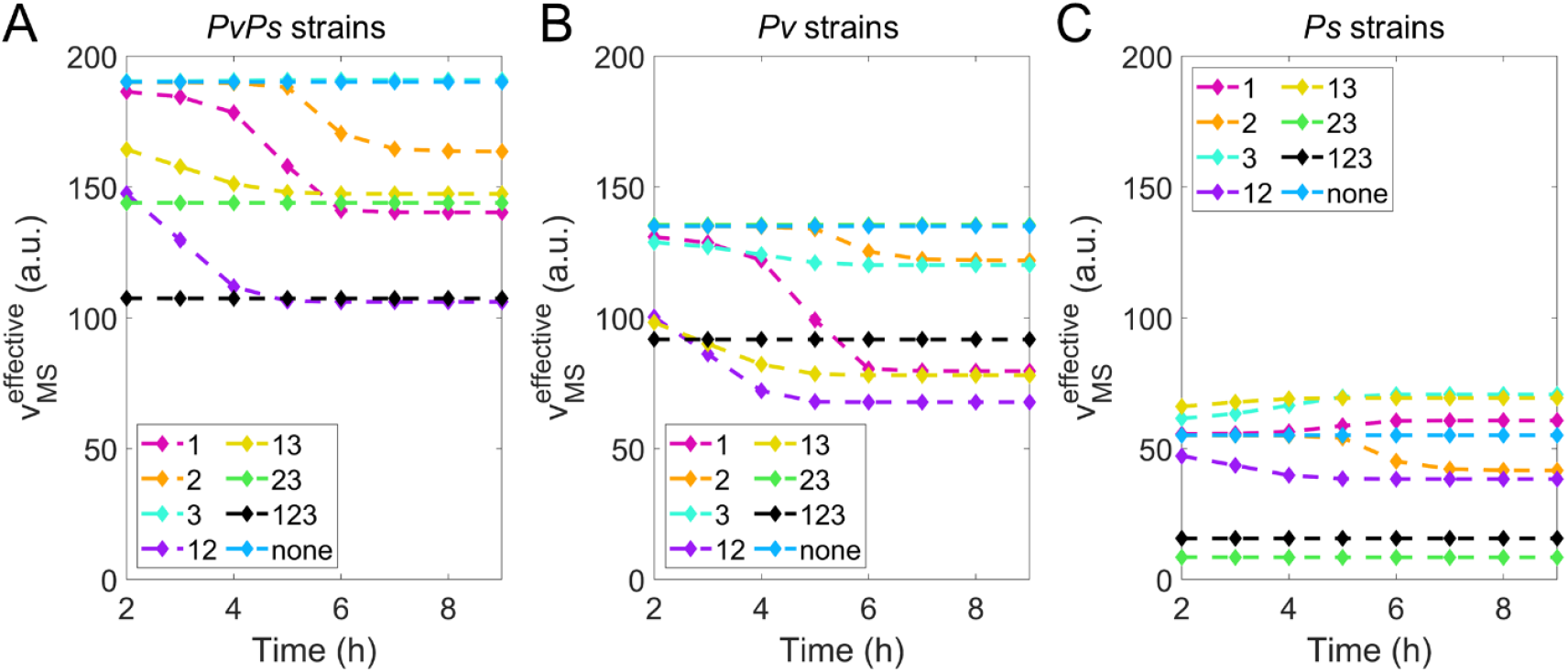
Model-predicted gene expression for all mutant strains computed with constant sigma factor levels. Each plot corresponds to a promoter-specific strain **A)** ‘P*v*P*s* strains’ with constant *σ*^*A*^-RNAP and constant *σ*^*H*^-RNAP, **B)** ‘P*v* strains’ with constant *σ*^*A*^-RNAP and **C)** ‘P*s* strains’ with constant *σ*^*H*^- RNAP.

**Figure S8.**
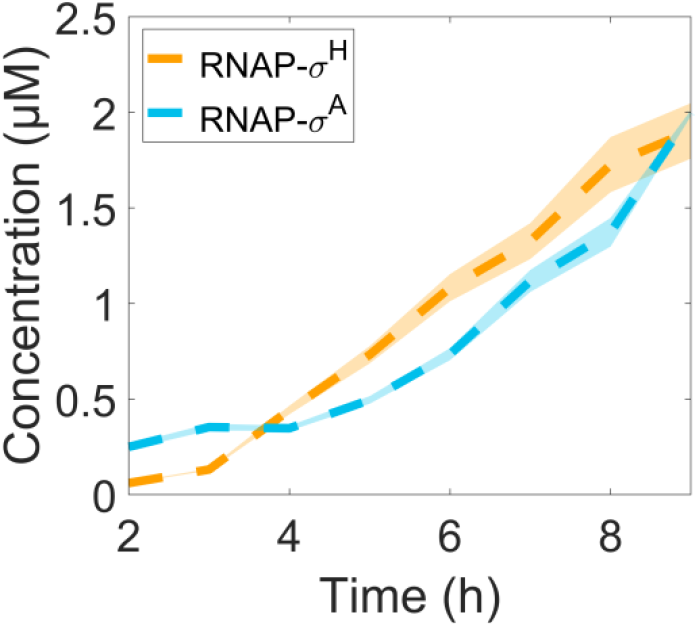
Model-predicted RNAP holoenzyme dynamics. Dashed lines are the mean concentration of RNAP holoenzyme bearing *σ*^A^ (blue) and *σ*^H^ (yellow). Colored patches are the standard deviations in predicted dynamics across 20 independent optimizations based on Table S4.

**Figure S9.**
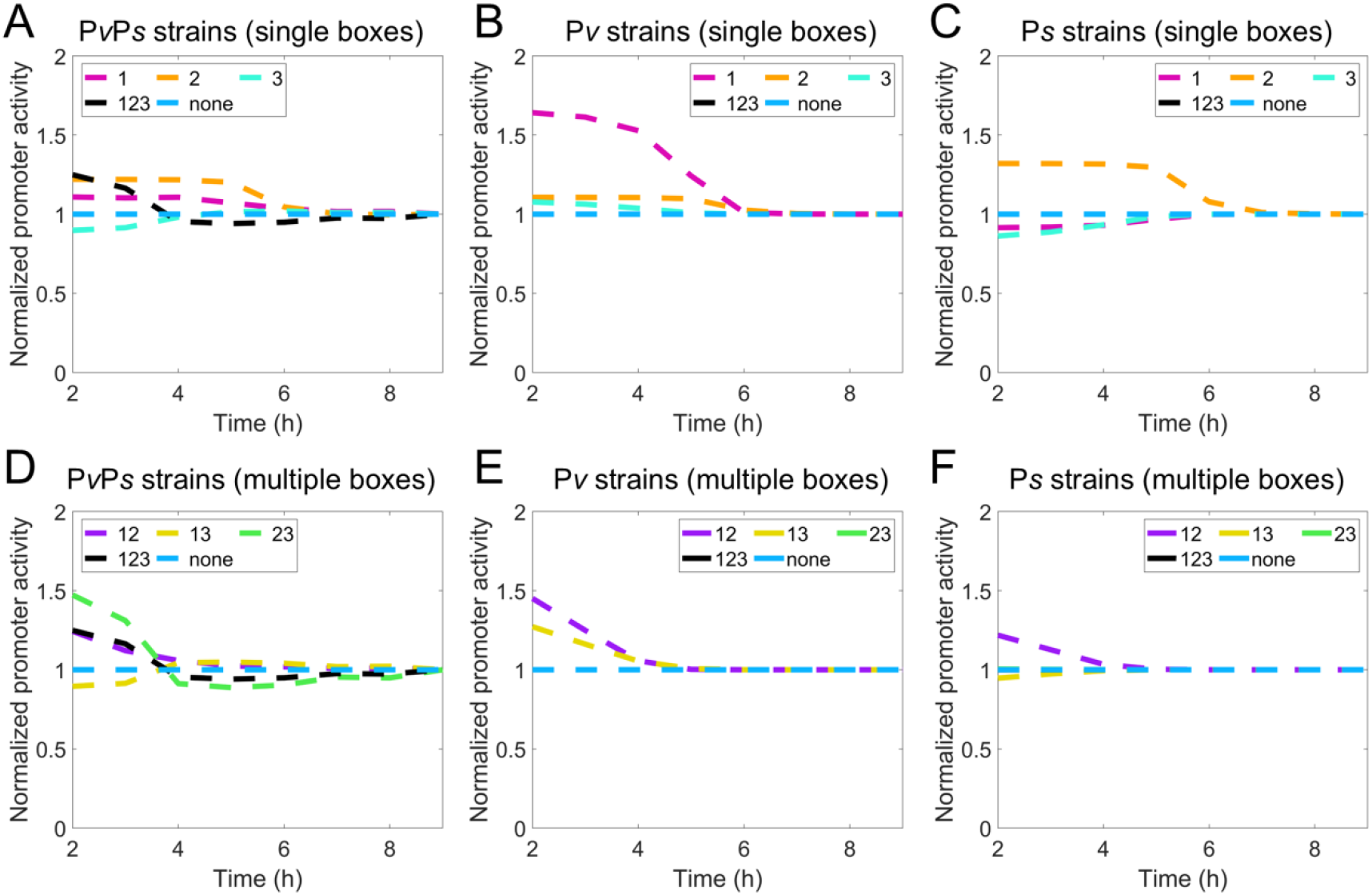
Normalized model-predicted effective transcription rate. Shown are normalized *v*^*eff*^ for strains with **(A-C)** single boxes and with **(D-F)** multiple boxes. Note the normalization is done by dividing the time-dependent prediction by its own final-time prediction, then by the prediction of ‘none’ for each promoter-specific construct. Normalization is performed for all dashed lines in Figure 5B-G.

**Figure S10.**
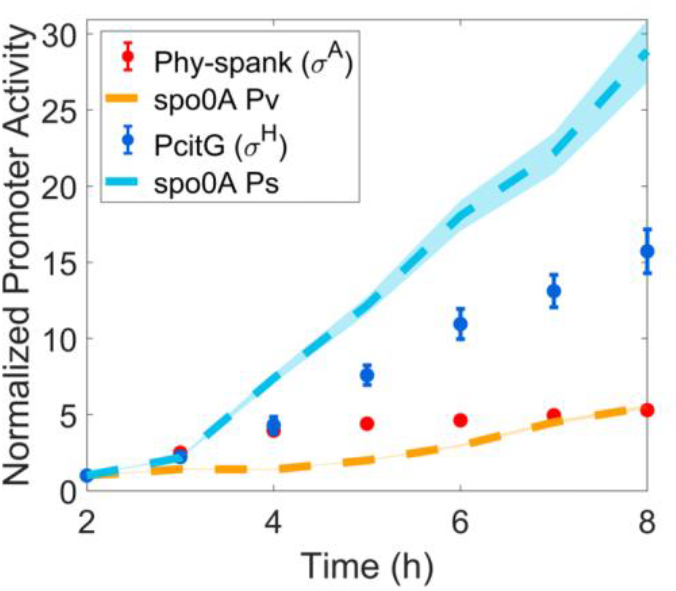
Sigma factor activities in *spo0A* promoter and other promoters. Normalized expression of two *σ*^A^-dependent promoter over time, *hyper-spac* and P*v* of *spo0A*; also shown is the normalized expression of two *σ*^H^-dependent promoters over time, *citG* and P*s* of *spo0A*. Note, data normalization is done by dividing each measurement by their corresponding measurement at t = 2 hours. Expression of *spo0A* P*v* and *spo0A* P*s* is the same as ‘P*v* strain, none’ and ‘P*s* strain, none’, respectively. The non-overlapping between blue dashed line and blue filled circles indicates *hyper-spac* and P*v* have different promoter binding affinity. Same is true for *citG* and P*s*.

**Figure S11.**
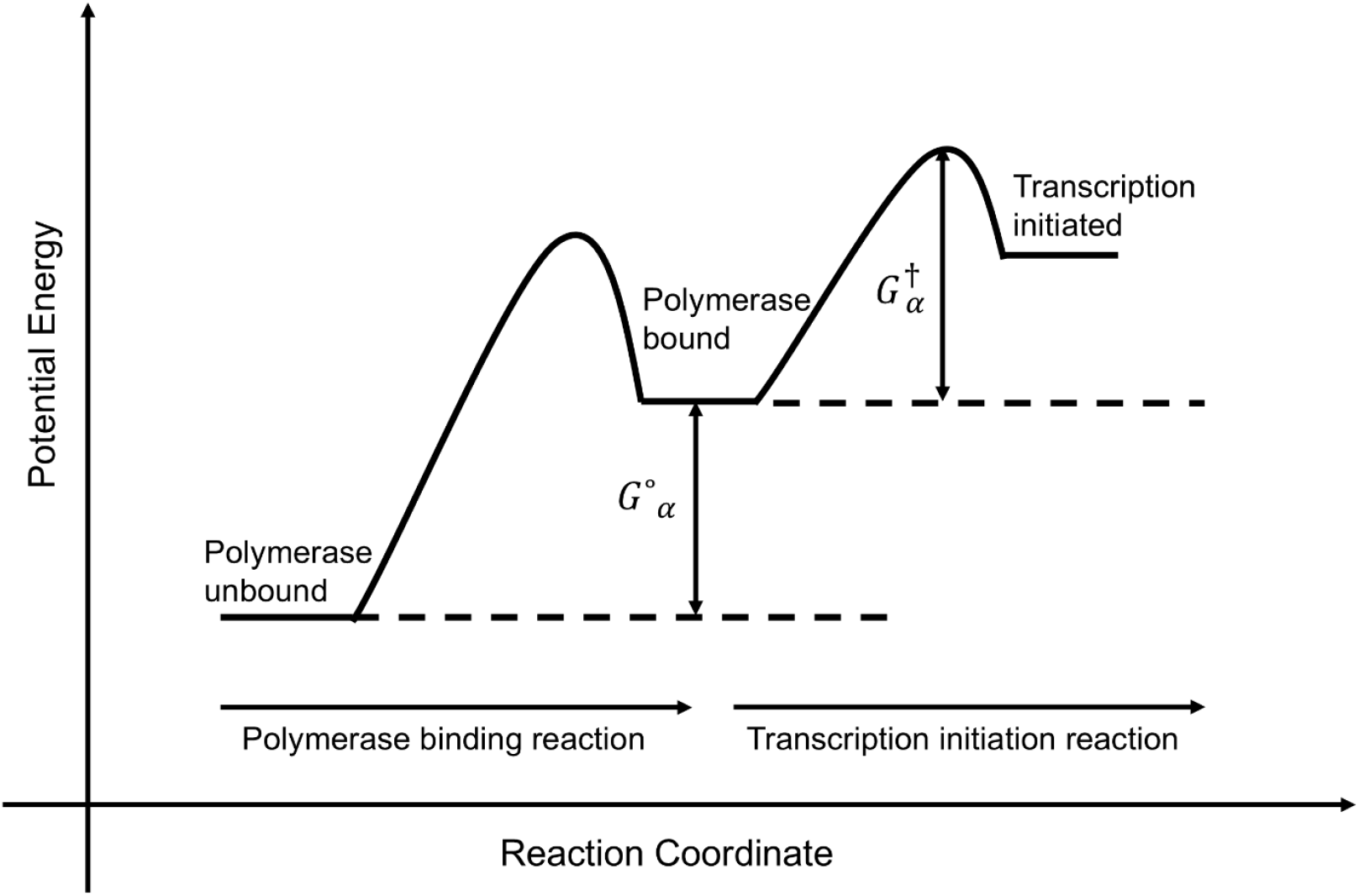
Energy diagram illustration of transcription, modeled as a two-step process. Reaction 1 corresponds to RNAP binding to the promoter (closed-complex formation). Reaction 2 corresponds to the post-closed-complex step. Namely, isomerization into an open complex and RNA synthesis initiation (8). Activation energy, *G*°_*α*_, between product and reactant state of reaction 1 determines the probability of RNAP binding (*Eqn. II. 4*). Activation energy of reaction 2, 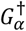, determines the maximum rate of transcription 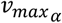 (*Eqn. II. 10*).

**Figure S12.**
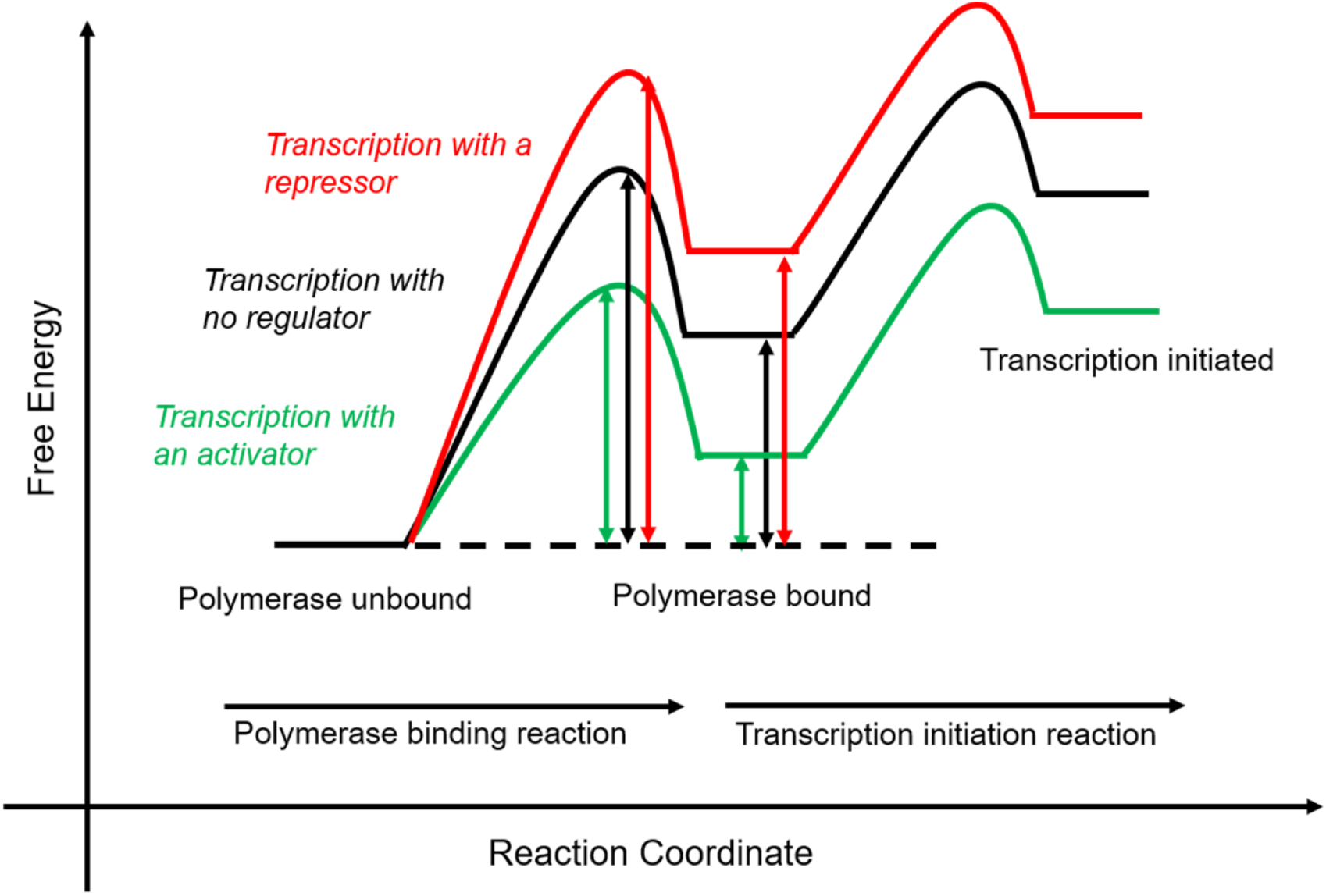
Energy diagrams for transcription with purely thermodynamic control. Upon the binding of transcriptional factor, an activator (repressor) lowers (increases) the potential energy for the polymerase binding reaction, indirectly lowering (increasing) the activation barrier in the same reaction, but there is no change in activation energy of the transcription initiation reaction.

**Figure S13.**
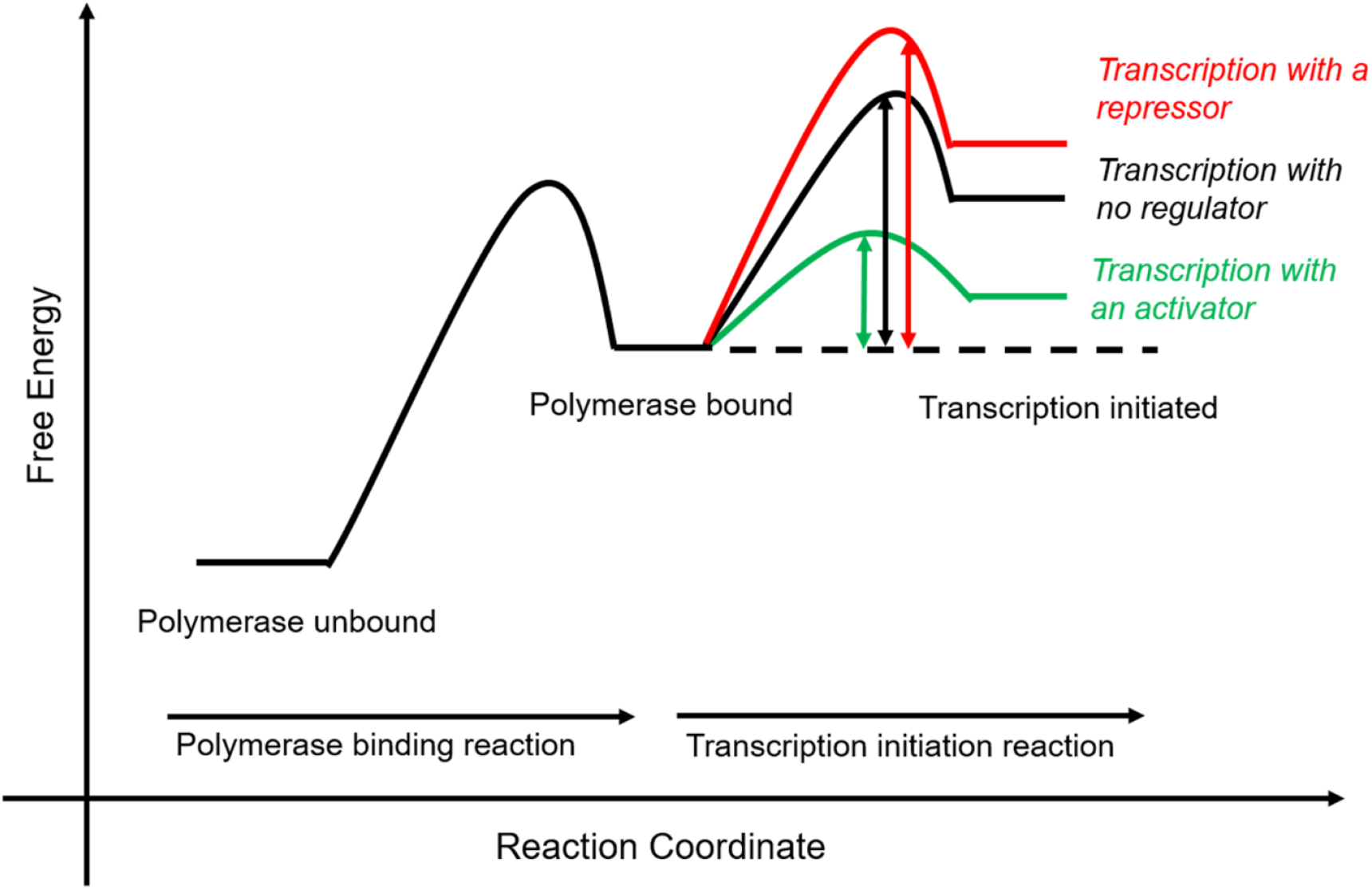
Energy diagrams for transcription with purely kinetic control. Upon the binding of transcriptional factor, an activator (repressor) lowers (increases) the activation energy for the transcription initiation reaction, but there is no change in the change in free energy of the polymerase binding reaction.

